# Perceptual repulsion and decisional attraction in male body serial dependence: a dimensional dissociation in channel architecture

**DOI:** 10.64898/2026.10.15.756928

**Authors:** John Cass, Daniel Talbot

**Author notes:** Corresponding author. School of Psychology, Western Sydney University, 74 Rickard Road, Bankstown, New South Wales, Australia. 2200.

## Abstract

Our judgments of the world are shaped by what we have just seen. This serial dependence is often treated as one bias, but it can constitute opposing influences: an attractive pull toward recent input and a repulsive push away from it. These have been linked to different stages, decisional and perceptual, but separated only for simple visual features. We asked whether the same separation holds for a socially significant category, the human body. Observers categorised images of male bodies as low or high on body fat and muscularity (binary choice, N = 74); a trial-level analysis estimated the perceptual and decisional contribution to each judgment. For body fat, the perceptual contribution was repulsive and clearly tuned, following a derivative-of-Gaussian profile peaking at a moderate difference between successive bodies (σ ≈ 5.6 standardised units), while the decisional contribution was attractive; the dissociation held in most observers. Muscularity showed the same attractive decisional bias, but its perceptual tuning emerged only at longer viewing times, where its width approached that for body fat, indicating a difference in resolvability rather than an absence of tuning. To our knowledge, this is the first estimate of the tuning of the perceptual repulsion channels from binary categorisation of continuously varying stimuli, the first separation of perceptual and decisional components in body perception, and the first evidence of serial dependence for male bodies. Perception emphasises how a body differs from the previous one while the decision favours continuity, a division of labour spanning low-level features to high-level body judgments.

## Introduction

Body misperception is a defining feature of several psychiatric conditions, most notably anorexia nervosa, and body dysmorphic disorder, including its muscularity-focused presentation known as muscle dysmorphia. People with anorexia nervosa systematically overestimate their own body size, even when the stimuli they judge are bodies of others (Glashouwer et al., 2019). People with muscle dysmorphia, by contrast, perceive themselves as smaller and less muscular than they objectively are (Tod et al., 2016). Even in non-clinical populations, body-composition judgments are systematically biased: thin bodies tend to be overestimated in size, and overweight bodies underestimated (Cornelissen et al., 2015; Cornelissen et al., 2017). The mechanisms underlying these biases, and their relationship to attitudinal and socio-cognitive factors with which they often correlate, are not fully understood. The dominant cognitive-behavioural account emphasises socio-cognitive contributors such as internalisation of media body ideals, social comparison, and dissatisfaction with one’s body (Tylka, 2011; van den Berg et al., 2002). However, an emerging line of work demonstrates that *purely perceptual* mechanisms (operating without reference to attitudes, beliefs, or social meaning) contribute substantially to body misperception (reviewed in (Brooks, Mond, et al., 2020). The present study contributes to this perceptual-mechanism line of work.

### Visual adaptation as a model of body misperception

The most influential perceptual account of body misperception treats body composition as encoded in the visual system via tuned neural channels, in the same way that orientation, motion direction, gaze direction, and faces are channel-coded (e.g., (Clifford, 2002; Webster & MacLin, 1999). According to this account, prolonged exposure to bodies at extreme positions along a body-composition dimension selectively adapts the channels tuned to that body type, producing a measurable shift in subsequent perception in the opposite direction: a classical visual aftereffect. Brooks and colleagues have established this account for body composition through a now-extensive programme of psychophysical work (Brooks, Keen, et al., 2020; Brooks, Mond, et al., 2020; Stephen et al., 2018; Sturman et al., 2017).

A key finding from this programme is the demonstration that body fat and muscularity adapt independently. In Sturman et al. (2017), participants viewed adapting bodies that varied in either body fat (low or high) or muscle mass (low or high) and then adjusted a test body to appear “normal” using orthogonal controls for each composition dimension. Adaptation produced aftereffects that were *dimension-specific*: adaptation to high-fat bodies shifted the normality setting on the fat axis but not on the muscle axis, and vice versa. The authors concluded that “adaptation can occur independently along the different dimensions of body composition” and that “neural mechanisms” are “independently sensitive to fat and muscle” (Sturman et al., 2017, p. 7). Brooks, Keen, et al. (2020) extended this work by demonstrating that the body-composition aftereffects also depend systematically on the gender of both the adapting and test bodies, indicating that the underlying channels carry rich representational structure beyond body composition alone. The broader Brooks, Mond, et al. (2020) review consolidates the channel-based account as the working framework for body-perception aftereffect research.

These adaptation studies establish that body perception is mediated by tuned neural mechanisms that are at least partially channel-specific. They leave open important questions, however, about the temporal dynamics of these mechanisms. Adaptation paradigms typically use prolonged exposure to a single (or small set of) adapting stimulus and measure the resulting aftereffect on a single subsequent test stimulus. This is well suited to characterising the architecture of the visual system at the relatively slow timescale of seconds to minutes, but cannot easily capture the moment-to-moment biases that arise when observers make many sequential judgments under naturalistic viewing conditions. To examine such effects we turn to a complementary paradigm: serial dependence.

### Serial dependence: trial-by-trial biases in perceptual decisions

Serial dependence is the phenomenon that perceptual judgments are systematically biased toward recently-viewed stimuli on a trial-by-trial timescale (Fischer & Whitney, 2014) reviewed in (Cicchini et al., 2024; Manassi et al., 2023). Since the seminal demonstrations in 2013 (Cicchini et al., 2014; Fischer & Whitney, 2014; Liberman et al., 2014; Van der Burg et al., 2013), serial dependence has been observed across an essentially open-ended set of feature dimensions: spatial position (Manassi et al., 2018), motion direction (Alais et al., 2017), numerosity (Cicchini et al., 2014), facial identity, expression, and attractiveness (Liberman et al., 2014; Liberman et al., 2018; Taubert et al., 2016), and gaze direction (Alais et al., 2018), among others. The phenomenon has also been demonstrated cross-modally and in the temporal domain (Van der Burg et al., 2013; Van der Burg & Goodbourn, 2015).

Two empirical signatures of serial dependence have become methodological foundations in the literature. The first is its featural-distance tuning: serial dependence is typically maximal when the inducer (previous trial) and target (current trial) are moderately similar, and decays for very similar or very dissimilar pairings. The shape of this distance-tuning is generally well captured by a derivative-of-Gaussian (DoG) function (Fischer & Whitney, 2014), which is also the analytic signature predicted by channel-based encoding accounts (in which the bias arises from a population of tuned channels overlapping at intermediate distances). The DoG shape, and its bandwidth parameter, are therefore not merely empirical descriptors but theoretically informative observables: a DoG-shaped perceptual bias is a candidate fingerprint of tuned-channel architecture (see also (Cicchini et al., 2017). The second signature is the relationship between serial dependence magnitude and stimulus reliability: less reliable target stimuli elicit larger serial-dependence effects (Ceylan et al., 2021; Cicchini et al., 2018; Gallagher & Benton, 2022). This second observation has been formalised as a Bayesian / Kalman-filter ideal-observer account (Burr & Cicchini, 2014; Cicchini et al., 2018; van Bergen & Jehee, 2019), in which the optimal weight given to the prior stimulus scales with the relative reliability of current and prior input.

On the face of it, serial dependence and adaptation seem to oppose one another. Adaptation generally produces a *repulsive* aftereffect (perception shifts *away* from the adapting stimulus) (Clifford, 2002), whereas serial dependence at the marginal level produces an *attractive* bias (perception shifts *toward* the previous stimulus) (Fischer & Whitney, 2014). The two effects can be observed simultaneously in the same experimental design (Pascucci et al., 2019), at different timescales, and with different signatures. Reconciling adaptation and serial dependence within a single computational framework is an active area of theoretical work (Fritsche et al., 2020; Pascucci et al., 2019; Sheehan & Serences, 2022) and is taken up further below.

### Serial dependence for body stimuli

Serial dependence has been documented in body perception in a series of studies, all of which have used reproduction measures (Alexi et al., 2018; Alexi et al., 2019; Turnbull et al., 2022). In reproduction tasks, observers do not classify the stimulus into a discrete category (such as “skinny” vs. “fat”); rather they produce a graded estimate of the perceived stimulus value, typically by positioning a marker on an analogue scale or adjusting a probe stimulus to match the target. (Alexi et al., 2018) introduced the most widely-used of these procedures for body perception, the “bodyline” task, in which observers position a stimulus body along an analogue line delineated by extreme body sizes. Using this task, Alexi et al. demonstrated that judgments of female body size are systematically attracted toward the previously-viewed body. The effect was well captured by a Kalman-filter ideal observer (in the tradition of (Cicchini et al., 2014), and the magnitude of bias scaled with discrimination thresholds. Alexi et al. (2019) replicated the effect using computer-generated female body stimuli, demonstrating that serial dependence does not depend on the realism of the body images. Alexi et al. (2019b) reported a significant positive correlation between body-size serial dependence and Eating Disorder Examination Questionnaire scores in young women, suggesting that the magnitude of the perceptual bias is clinically informative. Turnbull et al. (2022) extended this work to ensemble body-size judgments, showing that body serial dependence interacts with the spatial integration of multiple bodies in the visual field.

The existing body serial-dependence literature thus establishes the phenomenon as robust, clinically informative, and applicable across naturalistic and synthetic body images. Several gaps remain. First, all of this work has exclusively used female body stimuli; serial dependence for *male* body perception has not been examined. Second, all of this work has measured body size (effectively, the body fat / adiposity dimension). Muscularity as a discriminable dimension has not been examined despite its central importance to male body image disturbance (Murray, 2017). Third, all of this work has used reproduction (“bodyline”) methodology rather than two-alternative forced choice (2AFC) discrimination, leaving open whether the effect manifests in the more analytically constrained 2AFC setting. Fourth, and most importantly for the present study, none of this work has explicitly decomposed the marginal serial-dependence pattern into the perceptual and decisional components that have become central to the broader serial-dependence theoretical literature.

### Perceptual and decisional components of serial dependence

A foundational question in the serial-dependence literature concerns where the bias arises. If the bias originates at perceptual processing (i.e., if the previous stimulus actually alters the appearance of the current stimulus), then serial dependence is genuinely a perceptual phenomenon. If the bias arises post-perceptually (at decision, working memory, or response selection), then serial dependence is a decisional phenomenon that gives a misleading impression of altered perception. This is not a merely terminological distinction: the two accounts make different predictions about the neural locus of the effect, its temporal dynamics, its sensitivity to attentional and decisional manipulations, its relationship to other perceptual phenomena (notably adaptation), and its clinical interpretation. An influential early body of evidence in the orientation domain pointed in different directions, with (Fischer & Whitney, 2014) and (Cicchini et al., 2017) emphasising perceptual origins, and other lines of evidence pointing toward decision-stage or working-memory mechanisms (Bliss et al., 2017; Kiyonaga et al., 2017).

The first compelling within-paradigm dissociation was provided by Fritsche et al. (2017). They contrasted two complementary measurement procedures within the same observers: a stimulus reproduction task (in which the observer reproduces the orientation of the test stimulus after a brief delay), and a direct comparison task (in which the test stimulus was judged against a simultaneously-visible reference stimulus, minimising the contribution of working memory and decisional inertia). The stimulus reproduction procedure produced the canonical attractive serial-dependence pattern, but the direct comparison procedure revealed a *repulsive* effect of previous stimuli on the current judgment, particularly when the previous stimulus appeared at the same spatial location as the current one. Pascucci et al. (2019) formalised this dissociation through multilevel modelling and cross-validation, demonstrating that the marginal pattern reflects two opposing processes operating at distinct stages of processing: a stimulus-driven repulsion (related to neural adaptation) and a decision-driven attraction (related to decisional inertia). They proposed that the marginal serial-dependence pattern is the algebraic *sum* of these two components, with attractive serial dependence emerging where decisional inertia dominates and repulsive adaptation emerging where it does not. This two-process account has since been extended and corroborated in several directions. Gallagher and Benton (2022); (Gallagher & Benton, 2024) demonstrated that the apparent perceptual modulation of serial dependence by stimulus uncertainty is consistent with a decisional-attraction framework, and explicitly characterised serial dependence as predominantly an attraction to prior response rather than to prior stimulus. Ceylan et al. (2021) and Pascucci and Plomp (2021) used ‘no-report’ paradigms to dissociate the two components experimentally, confirming that perceptual repulsion and decisional attraction can be elicited independently.

The present analyses build on a methodological development from our own group (Cass et al., 2026) who recently developed a generalised linear mixed model (GLMM) approach to the perceptual/decisional decomposition specifically for 2AFC data. By including both the previous-stimulus and the previous-response covariates in the trial-level model, the partial-regression coefficient on each isolates the contribution of one source while controlling for the other. The decomposition can thus be conducted directly on the 2AFC psychometric function, removing the need for reproduction data and allowing application at the trial-by-trial timescale across long experimental sessions. To our knowledge, this combination of methodological features (2AFC binary choice, trial-level mixed-effects logistic regression, and simultaneous partialling of previous-stimulus and previous-response covariates) is new, although closely-related multiple-regression decompositions have been applied to reproduction and estimation data (e.g., (Pascucci et al., 2019). The present study is the first application of this framework to body perception, and, more broadly, the first application of a perceptual-decisional serial-dependence decomposition to psychometric-function data derived from a 2AFC categorisation task.

### The channel signature in the perceptual component

The perceptual/decisional decomposition offers a further analytical advantage that is particularly relevant in the present context. If a stimulus dimension is mediated by tuned neural channels, as (Sturman et al., 2017) and (Brooks, Keen, et al., 2020) have demonstrated for body fat and muscularity in using a prolonged adaptation paradigm, then the perceptual residual, after decisional carryover has been partialled out, should exhibit the DoG signature predicted by channel-based encoding. Specifically, the magnitude of the repulsive perceptual bias should grow with the inducer–target featural distance up to approximately the channel half-width, then decay back to zero for inducers beyond the channel’s featural ‘receptive field’ range. The DoG-shape signature has been treated as the empirical fingerprint of channel-based perception in many domains, including orientation (Fischer & Whitney, 2014; Pascucci et al., 2019), faces (Liberman et al., 2014), gaze (Alais et al., 2018), and position (Manassi et al., 2018). Whether body composition exhibits the DoG signature when measured via the perceptual residual, rather than via the marginal pattern (which is contaminated by decisional carryover), has not, to our knowledge, been tested.

There is in fact a specific reason to expect the marginal pattern to misrepresent the underlying channel architecture in 2AFC body discrimination. As noted, the marginal pattern is the sum of a broad perceptual repulsion and a decisional attraction (Pascucci et al., 2019). The decisional attraction itself peaks at large inducer-target differences (where the previous response is most likely to differ from the current response) and contributes a strong attractive shoulder to the marginal pattern. The interaction of broad perceptual repulsion and large decisional attraction generates a marginal DoG that is artefactually narrow: it appears to reveal channels with bandwidths far smaller than those that actually mediate perception. This is a generic consequence of partialling and was emphasised by (Pascucci et al., 2019). The partialled perceptual residual avoids this contamination and is therefore the appropriate analytical target for testing channel-based encoding accounts.

### The present study

We applied the trial-level GLMM partialling framework of (Cass et al., 2026) to a 2AFC body-discrimination dataset with three specific objectives. First, we asked whether the perceptual/decisional decomposition established for orientation (Fritsche et al., 2017; Pascucci et al., 2019) generalises to body perception, and whether the marginal pattern is correctly recovered as the predicted sum of the two components, providing a quantitative empirical test of the central theoretical claim of (Pascucci et al., 2019). Second, we asked whether each of two orthogonal body-composition dimensions, adiposity (body fat) and muscularity, exhibits the DoG signature of tuned-channel encoding in the perceptual residual. Given the prior demonstration that adaptation to both dimensions is dimension-specific (Brooks, Keen, et al., 2020; Sturman et al., 2017), the channel-based account predicts DoG signatures in both dimensions. Third, we asked whether individual differences in discrimination ability (the just-noticeable difference, JND) moderate any channel signature found, since both the Bayesian/Kalman-filter account (Burr & Cicchini, 2014; Cicchini et al., 2018) and the channel-based account make specific predictions about how individual differences should manifest.

Our dataset comprises male participants, each completing a single-image 2AFC body-discrimination task across two task blocks, one targeting body fat and one targeting muscularity. Two features of this design extend the prior body serial-dependence literature: it samples male observers (rather than the female samples used by (Alexi et al., 2018; Alexi et al., 2019; Turnbull et al., 2022), and it includes muscularity as a fully crossed discriminable dimension alongside body fat.

If serial dependence in body perception follows the same two-component structure established for simpler visual features, the partialled perceptual residual should be repulsive (the current body appearing pushed away from the preceding one) and the decisional carryover should be attractive (a tendency to repeat the preceding categorical response). Neither has been demonstrated for bodies before. Beyond this, if the perceptual component reflects adaptation in dimension-specific channels, as the prolonged-adaptation literature implies, the perceptual residual should carry a DoG-shaped tuning signature whose bandwidth reflects the width of those channels, and this signature should be estimable from the partialled residual in a way that the raw marginal pattern does not permit. What we add is a within-subject test of these predictions across two orthogonal body-composition dimensions, using the partialled perceptual residual as the measure, in a 2AFC task that for the first time permits the channel-tuning profile to be estimated rather than assumed.

## Method

The behavioural dataset analysed here was collected as part of a study reported by (D. Talbot et al., 2019), where the apparatus, stimuli, and body-categorisation procedure are described in full. That earlier paper related summary psychophysical indices (point of subjective equivalence, just-noticeable difference, and reaction time) to self-report measures of body dissatisfaction and eating-disorder symptoms. Our study re-analyses the same trial-level data with an entirely different aim and entirely new methods: the trial-by-trial decomposition of serial dependence into perceptual and decisional components, and the estimation of perceptual channel tuning. None of the serial-dependence analyses, the perceptual/decisional decomposition, the channel-tuning estimation, or the simulations reported here appear in the earlier paper or anywhere else, only the raw categorisation data are shared between the two studies. We summarise the design below at the level needed to interpret the present analyses, and refer the reader to (D. Talbot et al., 2019) for the complete procedural detail.

### Participants

Eighty-two male undergraduate students from Western Sydney University took part in the original study (age range 17 to 35 years, M = 21.89, SD = 4.20; (D. Talbot et al., 2019)). The sample size was set a priori by a power analysis for the correlational aims of that study (G*Power; α = 0.05, β = 0.20; medium effect, ρ = 0.30, two-tailed), which indicated 82 participants. All procedures were approved by the Western Sydney University Human Research Ethics Committee (ethics ID: H11778; approved 22 August 2016) and were conducted in accordance with the Declaration of Helsinki. All participants provided informed consent prior to participation.

For the present trial-level analyses, a participant was retained only if the body-categorisation task yielded usable trial-by-trial data in both the body fat and muscularity blocks. Eight of the 82 participants were excluded on this criterion: for seven (participants 2, 12, 28, 31, 32, 40, and 44) the stored data contained only figure-rating-scale summary values and no trial-level categorisation records, and for one (participant 14) the trial file was malformed and could not be parsed. None of the eight contributed any categorisation trials, so no partial data were discarded. The final sample comprised 74 participants, each with complete data in both blocks, yielding 35,520 body fat trials and 35,531 muscularity trials (71,051 trials in total).

### Stimuli

Body stimuli were computer-rendered images of male bodies varying along two independent dimensions, body fat and muscularity, generated as described in (D. Talbot et al., 2019). The stimulus sets were derived from the Visual Body Scale for Men (Daniel Talbot, John Cass, et al., 2019) and the New Somatomorphic Matrix-Male (Daniel Talbot, Evelyn Smith, et al., 2019), which depicted male bodies spanning body fat values from 4% to 40% and fat-free muscle index values from 16.5 to 30 kg/m^2^. The body categorisation task comprised 16 graded stimulus levels for the body fat (skinny-fat) dimension and 16 graded stimulus levels for the muscularity (scrawny-muscular) dimension. Each dimension was sampled across many graded levels rather than the small number of fixed exemplars typical of figural rating scales, with stimulus magnitude indexed on a body-number scale spanning 10 to 100 (21 distinct levels were presented across the pooled data, with a nominal midpoint of 55). The body categorisation task comprised 16 graded stimulus levels for the body fat (skinny-fat) dimension and 16 graded stimulus levels for the muscularity (scrawny-muscular) dimension (Figure 1). This dense, continuous sampling of each dimension is what permits the perceptual carryover to be characterised as a function of inducer-target distance, and therefore the channel-tuning profile to be estimated.

**Figure 1.**
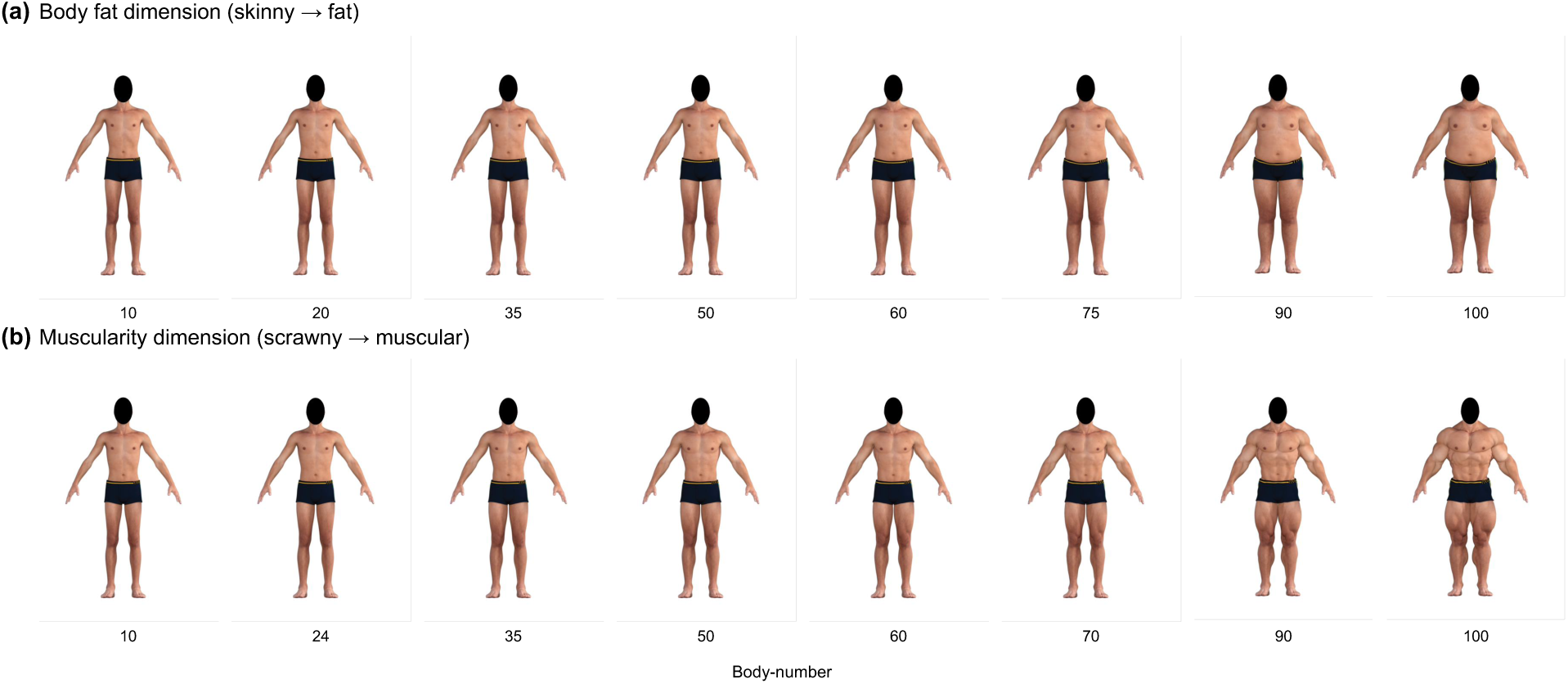
Example male body stimuli along the two judged dimensions. **(a)** The body fat dimension, from skinny to fat, with muscularity held at its reference value. **(b)** The muscularity dimension, from scrawny to muscular, with body fat held at its reference value. Each dimension comprised 16 graded levels indexed on a body-number scale from 10 to 100; eight evenly spaced levels are shown here, with the body-number beneath each image. Faces were masked and all images shared the same viewpoint, scale, and background. Renders were generated as described in Talbot et al. (2019).

### Design and procedure

Testing was conducted individually in a low-illumination laboratory to minimise visual distractions. Stimuli were presented on a 15.4-inch LED monitor (resolution 1280 × 800 pixels) at a fixed viewing distance of 340 mm. Participants first completed a practice block to familiarise themselves with the task before completing two experimental body-categorisation blocks. The order of the body fat and muscularity blocks was counterbalanced across participants to control for order effects. In the body fat block they categorised each body as “skinny” or “fat”; in the muscularity block they categorised each body as “scrawny” or “muscular”. These were single-stimulus judgements made against each participant’s own internal reference rather than against an externally presented standard. On each trial, a single body image was presented centrally on the screen and remained visible until the participant made a dichotomous keypress response. Following each response, the stimulus disappeared and a central fixation cross was presented for 500 ms before the next trial commenced. Reaction time was recorded from stimulus onset to the participant’s response. Each block comprised 16 graded stimulus levels spanning the relevant body-composition dimension. Each body image (166 pixels in height) was presented 30 times, yielding 480 trials in the body fat block and 480 trials in the muscularity block. Stimulus presentation order was randomised independently for each participant, such that body level varied unpredictably from trial to trial. This randomisation produced an approximately uniform distribution of inducer–target differences (Δ) across successive trials, providing the sampling density required for the present serial-dependence analyses. Because judgements were made relative to each participant’s own subjective criterion, the resulting point of subjective equality reflects an individual’s internal category boundary rather than an experimenter-defined or objectively veridical midpoint.

### Statistical analysis

All analyses were conducted in Python and R. Within each block, responses were coded dichotomously (1 = “fat” or “muscular”, 0 = “skinny” or “scrawny”), and stimulus magnitude (body-number) was z-scored within participant and block before analysis. The inducer-target distance on each trial was defined as Δ = stim(*t*−1) − stim(*t*) in z-units (one z-unit corresponds to approximately 10 body-number units). The immediately preceding trial (lag 1) was the primary inducer throughout. Lag was varied explicitly only in the temporal-decay analysis.

#### Marginal serial dependence

We first estimated the surface (marginal) influence of the previous stimulus on the current response, fitting a logistic model of the current response on the previous stimulus without controlling for the previous response. This provides the conventional, undecomposed serial-dependence estimate.

#### Perceptual/decisional decomposition

To separate the perceptual from the decisional carryover, we fitted a logistic generalised linear mixed model (GLMM) to the trial-level data within each block, predicting the current response from the current stimulus, the previous stimulus, and the previous response, with random intercepts for participants (and, where estimable, random slopes).

In this model the partial coefficient on the previous stimulus indexes the stimulus-driven (perceptual) carryover with the previous response held constant, and the partial coefficient on the previous response indexes the response-driven (decisional) carryover with the previous stimulus held constant. The perceptual residual referred to throughout is the current-response log-odds after the previous-response term has been partialled out; it is the quantity on which all channel-tuning analyses were performed. Where the full GLMM was computationally prohibitive (for example, across the many fits of the lag-decay analysis), an equivalent pooled logistic model with participant fixed effects was used.

#### Channel-tuning (DoG versus linear) model comparison

To test whether the perceptual residual carries the signature of tuned channels, we expressed it as a function of Δ and compared two nested models: a linear function (repulsion proportional to Δ, with no channel structure) and a derivative-of-Gaussian (DoG) function:

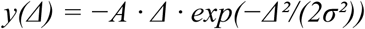

following the sign convention of (Fischer & Whitney, 2014), in which a positive amplitude A denotes repulsion and σ denotes the channel bandwidth. The two models were compared by likelihood-ratio test and by the difference in Akaike Information Criterion (ΔAIC). This comparison was performed at the population level (trials pooled across participants within each block) and, for the per-participant distributions, separately for each participant. Bandwidths that were not identifiable within a participant’s trial count were capped at an upper bound (reported where relevant).

#### Within-participant confirmation of curvature

Because a curved population-level function can in principle arise from averaging across participants who individually follow straight lines of differing slope, we verified that the body fat curvature was a genuine within-participant property. For each participant we partialled the linear trend out of the DoG-shaped predictor and estimated the coefficient on the orthogonal non-linear component. These per-participant coefficients were then submitted to a one-sample test against zero (t-test and Wilcoxon signed-rank). A reliably non-zero coefficient indicates curvature present within individuals rather than manufactured by pooling. We confirmed the same conclusion in a mixed model that allowed each participant their own intercept and linear slope.

#### JND-moderated tuning

To test whether discrimination ability moderates the tuning shape, participants were split at the median JND within each block and the distance-tuning function was re-estimated per subgroup. The dependence of tuning shape on dimension and discrimination ability was assessed by a three-way (task × JND subgroup × shape) likelihood-ratio test.

#### Individual differences

Per-participant DoG amplitude (A) and bandwidth (σ) were each correlated with per-participant JND, using Pearson and Spearman coefficients. JND was estimated from a logistic psychometric fit per participant and block, defined (following (D. Talbot et al., 2019), as half the difference between the stimulus levels yielding 75% and 25% “fat” or “muscular” responses.

#### Simulation

To test the claim that the narrow marginal DoG is the emergent sum of the broad perceptual and decisional components, we generated a synthetic dataset from the fitted joint-model components alone and re-estimated the marginal serial-dependence function from it, comparing its bandwidth and amplitude to the empirical marginal estimates.

#### Temporal decay

The persistence of each component was assessed by re-estimating the perceptual and decisional carryover at increasing lags between inducer and target, separately for each block.

Effects are reported with likelihood-ratio χ² and *p*-values for model comparisons, with ΔAIC where model preference is at issue, and with paired nonparametric tests (Wilcoxon) for within-participant contrasts. Exact test statistics and degrees of freedom are given with each result in the Results.

## Results

### Overview and analytic approach

After excluding eight participants whose stored files contained only figure-rating-scale summary values and no trial-level categorisation records (see Method), the analysed dataset comprised 71,040 trials in total. Trial-level analyses used mixed-effects logistic regression fitted with the lme4 package in R (Bates et al., 2015) with subject random intercepts. Individual-difference analyses used per-subject psychometric indices (point of subjective equality, PSE; just-noticeable difference, JND) derived from independent two-parameter logistic functions fitted to each participant’s data within each block, with the slope parameter (sensitivity) and PSE as free parameters. JND was computed as the half-distance on the stimulus axis corresponding to the 25– 75% response range. Per-subject regression coefficients were obtained from the same per-subject logistic fits and aggregated via paired t-tests, Wilcoxon signed-rank tests, and Pearson/Spearman correlations as appropriate. Correction was applied to each family of tests.

Our analytic framework is anchored in the perceptual/decisional decomposition of serial dependence pioneered by (Fritsche et al., 2017) and extended and formalised by (Pascucci et al., 2019). Both groups demonstrated, using reproduction orientation tasks, that the apparently attractive bias of serial dependence is the joint product of two opposing processes: a stimulus-driven repulsion (“vision goes for novelty”) and a response-driven attraction (“decisions for perseverance”). Our study extends this decomposition to 2AFC body discrimination data using a generalised linear mixed model (GLMM) framework (Cass et al., 2026) that partials out decisional carryover from stimulus-driven aftereffects at the trial level. We apply this framework to our two body-composition dimensions (adiposity and muscularity) within the same participants, allowing within-subject comparisons of perceptual and decisional architectures across dimensions. To our knowledge, serial dependence has not previously been measured in male body stimuli or in muscularity as a discriminable dimension; prior body-image serial-dependence work has used female body images and body size (e.g., (Alexi et al., 2018; Alexi et al., 2019; Turnbull et al., 2022).

We present results in a sequence that mirrors the logic of the analysis. We begin by showing the apparent absence of serial dependence at the marginal level. We then introduce the trial-level partialling and show that perceptual and decisional carryover components, individually masked at the marginal level, emerge robustly with opposite signs. We then demonstrate that the classic attractive “derivative-of-Gaussian” (DoG) serial-dependence tuning function is the predicted sum of these two opponent components. A series of diagnostic tests confirms that the decisional component is genuinely response-driven and that the stimulus-driven component is genuinely stimulus-driven. We then characterise the long-range temporal persistence of each component, establish that body fat (but not muscularity) shows the signature of tuned-channel encoding, and link the channel signature to individual differences in discrimination ability.

### Discrimination, bias, and sensitivity

Population psychometric functions for each block are shown in Figure 2. Observers reliably discriminated body composition in both dimensions: mean per-subject logistic slope coefficients for the current stimulus (*β_stim_t*, indexing sensitivity to the stimulus being judged on each trial) were β = 0.20 (body fat) and β = 0.15 (muscularity). Discrimination thresholds were modest: mean JND = 5.7 body-number units (SD = 3.1) for body fat and mean JND = 8.4 (SD = 3.0) for muscularity, with muscularity slightly less precise than body fat.

**Figure 2.**
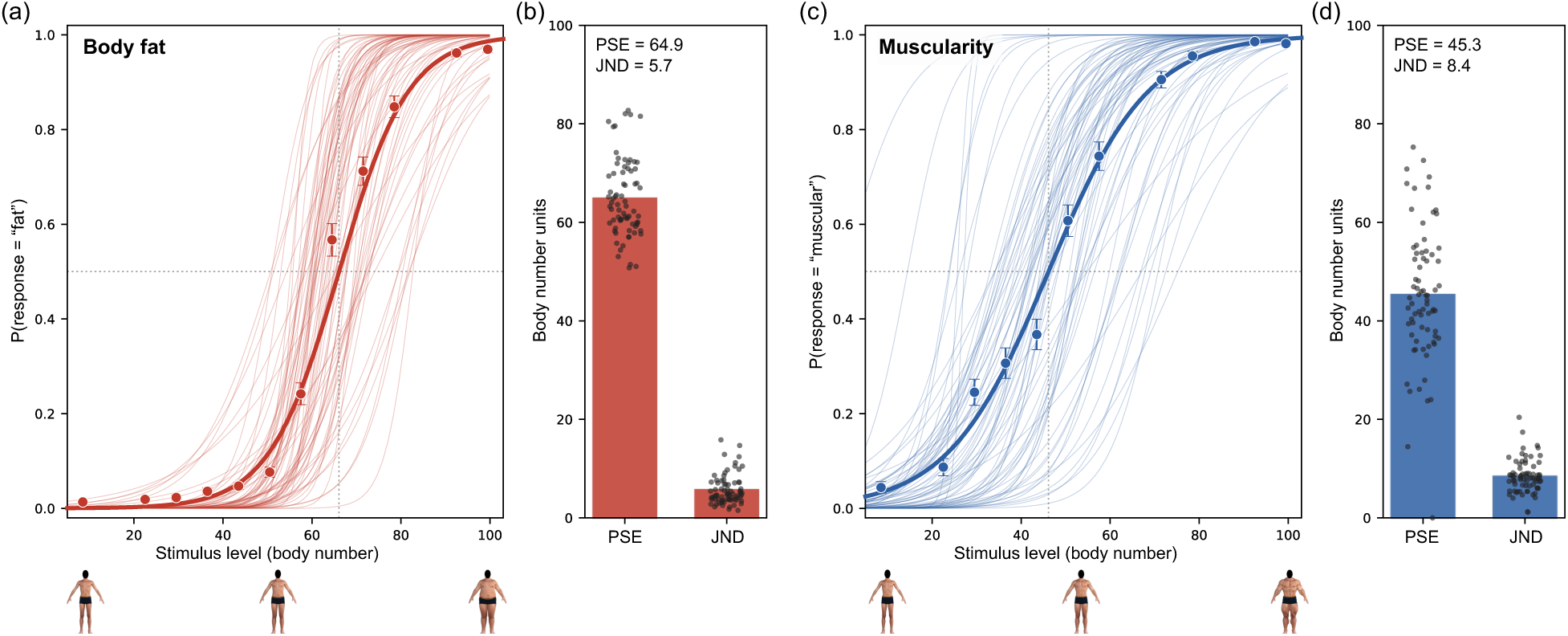
Marginal psychometric functions for body fat (a) and muscularity (c) blocks. Thin coloured lines show individual subjects (n = 74). Solid curves are logistic fits to the population-averaged data, not used in subsequent analyses. Markers and error bars show the population mean ± 95% CI. Side panels (b) and (d) show per-subject PSE and JND values (black dots) with population mean (bars).

The point of subjective equality (PSE) corresponds to the stimulus value at which an observer is equally likely to select either response option in a given block. Body number = 55, the midpoint of the 10–100 stimulus scale, represents an unbiased response criterion. PSEs deviated from this midpoint in opposite directions by block, indicating a conservative bias in body fat (mean PSE = 64.9, SD = 8.1; one-sample t(73) = 10.4, *p* < .001) and a liberal bias in muscularity (mean PSE = 45.3, SD = 13.3; t(73) = −6.2, p < .001). Observers were thus systematically reluctant to classify bodies as “fat” and equally willing to classify them as “muscular,” consistent with a population-level bias toward favourable body composition.

### Trial-level partialling unmasks two opposing components

We begin with the simplest model that could capture serial dependence at the trial level. It treats the current response as the outcome of a logistic regression with two stimulus-related predictors, the current stimulus and the previous-trial stimulus, plus a subject-specific baseline (a random intercept) that allows different observers to differ in their overall tendency to respond ‘high’ (that is, “fat” or “muscular”). We refer to this as the marginal model, because it does not separate perceptual from decisional carryover. The model is:

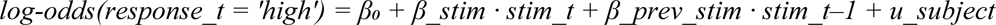

The log-odds of responding ‘high’ on trial t is the sum of a baseline (β₀), a current-stimulus contribution that scales with sensitivity (*β_stim*, larger values corresponding to steeper psychometric functions), a previous-stimulus contribution (*β_prev_stim*, the carryover term of interest), and a subject-specific offset (*u_subject*). A positive *β_prev_stim* would mean that a higher previous stimulus increases the log-odds of responding ‘high’ on the current trial (an attractive carryover); a negative *β_prev_stim* would indicate a repulsive carryover.

The marginal model returned a statistically reliable but small positive previous-stimulus coefficient in both blocks: *β_prev_stim* = +0.042 in body fat (SE = 0.010, z = 4.34, *p* < 10⁻⁵) and +0.082 in muscularity (SE = 0.010, z = 8.60, *p* < 10⁻¹⁷). These coefficients are 16 to 39 times smaller than the current-stimulus sensitivity coefficient (β_stim = +1.63 and +1.20, respectively), so previous-stimulus information contributed only marginally to current responses at the population level. Translated into PSE-shift units *(−β_prev_stim* / *β_stim*), the implied displacement of the psychometric function between the most extreme previous-stimulus conditions is approximately 1 body-number unit in body fat and 3 body-number units in muscularity, very small relative to the 90-unit stimulus span. Figure 3a and b show that at this magnitude the three previous-stimulus conditions yield closely overlapping psychometric curves, with the displacement barely discernible.

**Figure 3.**
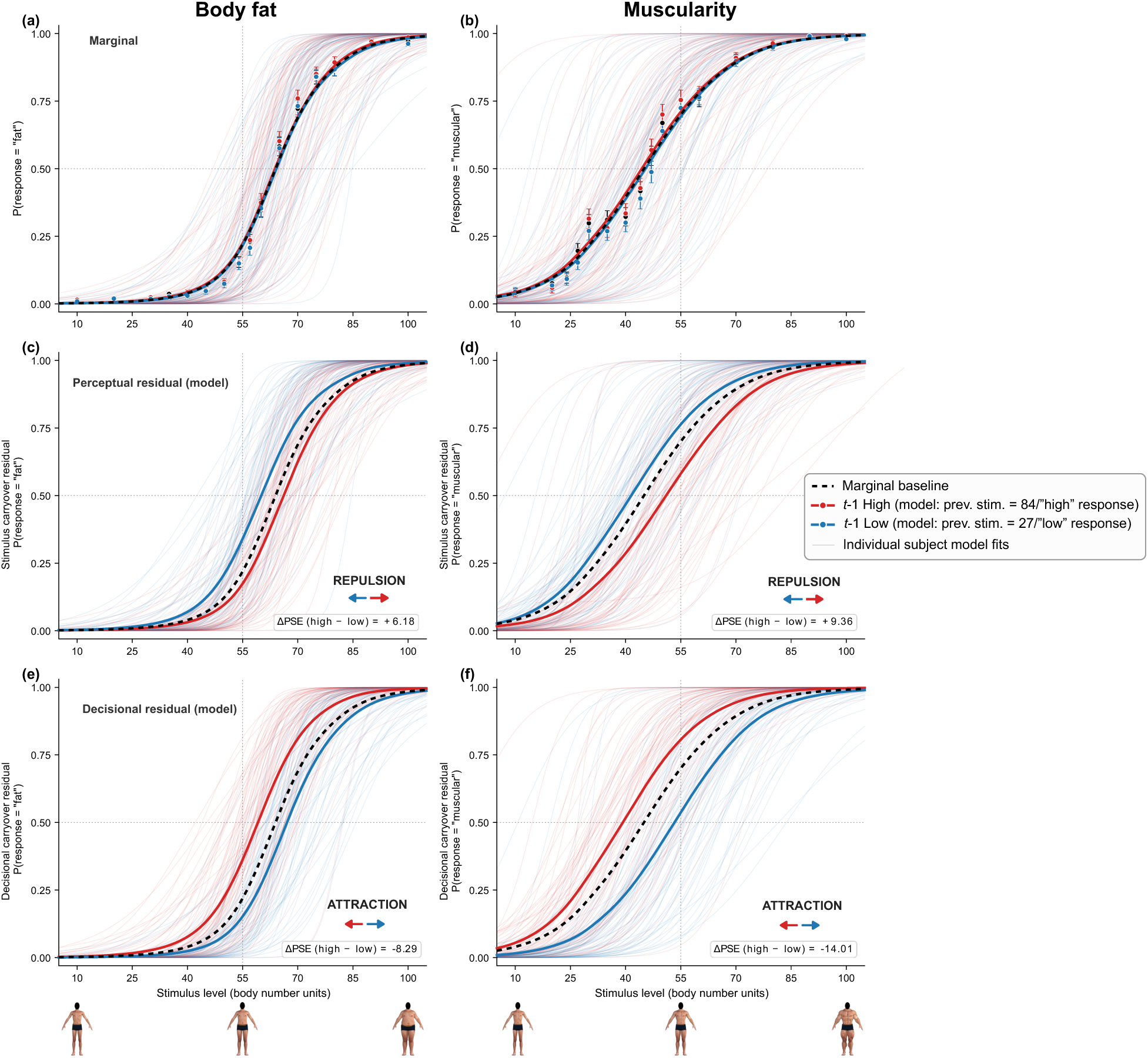
Marginal versus model-decomposed psychometric functions for body fat (left column) and muscularity (right column). **(a, b)** plotted separately for each previous-stimulus category, with no partialling. Markers show the between-subject mean P(response) ± 1 SEM. The three curves, one per previous-stimulus condition, are visually indistinguishable, consistent with an apparent absence of serial dependence at the marginal level. **(c, d)** The stimulus carryover (perceptual) component, taken from the joint trial-level logistic model (*response_t ∼ stim_t + prev_stim + prev_resp_centred*). Curves are model-predicted at a low (*prev_stim* = 27, blue) and a high (*prev_stim* = 84, red) previous stimulus, with the previous response held at its mean. The red curve shifts to the right of the dotted marginal reference and the blue curve to the left, the signature of perceptual repulsion: the current body is judged away from the previous one. **(e, f)** The decisional carryover component, from the same joint model. Curves are model-predicted at the two previous responses (±0.5), with the previous stimulus held at its mean. The red curve shifts to the left and the blue curve to the right, the signature of decisional attraction: responses are pulled toward the previous response. Thin transparent lines show per-subject model fits. The black dotted curve in each panel is the marginal psychometric function for the current trial, collapsed across previous-stimulus and previous-response conditions., included for visual reference. Coloured arrows mark the PSE shift of each conditional curve relative to the marginal, and inset values report the PSE shift between the red and blue curves in body-number units (positive = repulsion in c, d; negative = attraction in e, f).

Taken at face value, these marginal estimates suggest only a weak and, if anything, attractive influence of the previous stimulus on the current response, apparently inconsistent with the substantial attractive biases reported in earlier body-size work (Alexi et al., 2018; Alexi et al., 2019) and with the broader serial-dependence literature. As we show next, this small marginal pattern masks two much larger opposing effects.

To dissociate the perceptual and decisional contributions, we re-fitted the same data with a third predictor added, the previous-trial response, coded ±0.5 (’high’ = +0.5, ‘low’ = −0.5). The centring is on the response scale itself, not on the participant’s PSE or the stimulus axis, so that the coefficient on the previous response reads directly as the change in the log-odds of responding ‘high’ (“fat” or “muscular”) when the previous response was ‘high’ rather than ‘low’ (“thin”/“scrawny”). We refer to this as the joint model:

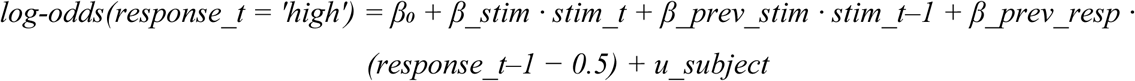

In other words, the joint model is the marginal model plus a decisional term that depends on what the observer responded on the previous trial. *β_prev_resp* is the decisional carryover coefficient, and a positive value indicates a tendency to repeat the prior response. For completeness, an intermediate model with only the previous response and not the previous stimulus is conceptually possible, but it would not isolate the perceptual component, because the previous response and the previous stimulus are correlated. The joint model estimates both terms simultaneously, and both partialled components are read from this single fit.

Adding the previous-response term produced a striking realignment of the previous-stimulus coefficient. In the body fat block, *β_prev_stim* switched sign from +0.042 (marginal model) to −0.215 (joint model; SE = 0.015, z = −14.82, *p* < 10⁻⁴⁹). The new coefficient indicates that, once decisional carryover is held constant, the previous stimulus exerts a repulsive influence on the current response: a higher previous stimulus reduces the log-odds of responding ‘high’ on the current trial. In the muscularity block, the same sign reversal occurred, from +0.082 to −0.249 (SE = 0.014, z = −18.21, *p* < 10⁻⁷³). Simultaneously, the previous-response coefficient emerged with a large positive value: *β_prev_resp* = +1.53 in body fat (SE = 0.065, z = 23.5) and +2.00 in muscularity (SE = 0.059, z = 34.1), both *p*-values < 10⁻¹⁰⁰, indicating a strong tendency to repeat the prior response.

Likelihood-ratio tests confirmed that both terms contributed independently and substantially: adding the previous-response term to the marginal model produced χ²(1) = 558 (body fat) and χ²(1) = 1297 (muscularity), and adding the previous-stimulus term to a decisional-only baseline produced χ²(1) = 238 (body fat) and χ²(1) = 506 (muscularity), all *p* < 10⁻⁵⁰. Per-subject fits confirmed that the sign-flip is not an aggregation artefact: 84% of subjects in the body fat task and 92% in the muscularity task individually showed a negative *β_prev_stim* once the decisional carryover was partialled out.

The implication is that the apparent near-absence of serial dependence at the marginal level (the small marginal coefficient above, and the overlapping psychometric curves in Figure 3a and b) reflects the near-cancellation of two large opposing influences: a perceptual residual that is repulsive (current judgments shift away from the previous body stimulus), and a decisional carryover that is attractive (current judgments are pulled toward the previous response). Figure 3c and d show that, once decisional carryover is partialled out, the three previously near-coincident psychometric curves (one per previous-stimulus condition) separate horizontally, with their PSEs displaced in the repulsive direction. Figure 3e and f show the matching decomposition for the decisional carryover, conditioned on the previous response: the curves shift in the attractive direction. Both effects are large, and the dissociation in direction is unambiguous.

The sign-flip we observe in the perceptual residual when decisional carryover is partialled out is a precise replication of the central finding of Fritsche et al. (2017) and Pascucci et al. (2019) in the orientation domain, now extended to two body-composition dimensions and to 2AFC body discrimination. The partialling framework therefore generalises to body perception, and the perceptual/decisional dissociation is preserved across both stimulus dimensions tested here.

### The classic attractive DoG serial-dependence shape is the predicted sum of the two components

A long-standing observation in the serial-dependence literature is that the classic attractive bias follows a derivative-of-Gaussian (DoG) tuning function: the current judgment is most strongly attracted to the previous stimulus at intermediate inducer-target distances, and the bias falls off at very small and very large distances (Fischer & Whitney, 2014). Let Δ denote the signed difference between the previous and current stimulus: Δ = stim *t*–1 – stim *t* (in z-units, where one z-unit corresponds to ten body-number units throughout). The DoG curve is:

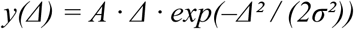

Here, y(Δ) is the magnitude of the serial-dependence bias when the previous stimulus differs from the current stimulus by Δ. The amplitude A indicates the direction and strength of the bias: positive A means the current response is biased toward the previous stimulus (attraction); negative A means biased away (repulsion). The bandwidth σ is the value of Δ at which the bias is maximal, and is conventionally interpreted as the half-width of the underlying perceptual channel. We use the standard sign convention (Fischer & Whitney, 2014) throughout.

Below we describe how the marginal DoG (Figure 4 c and d, solid line), the perceptual DoG (green curve, Figure 4 a and b), the decisional sigmoid (purple curve, Figure 4 a and b), and the simulation-recovered marginal DoG (Figure 4 c and d, dashed) are each obtained from the same trial-level dataset. The procedure is mechanically simple but each step matters for interpretation, so we walk through each in turn.

**Figure 4.**
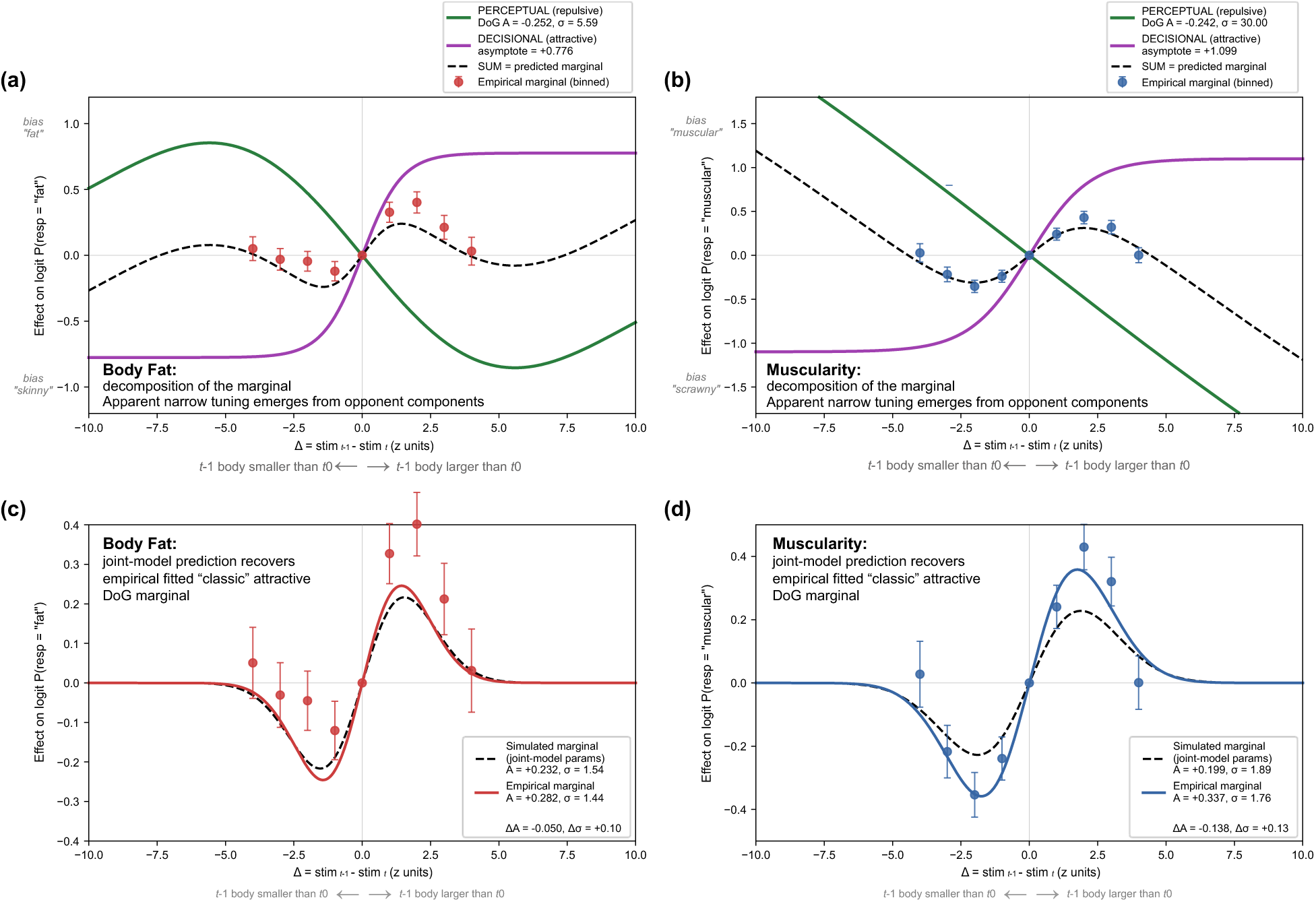
The classic attractive DoG serial-dependence shape is the predicted sum of perceptual repulsion and decisional attraction. Top row (panels a, b): decomposition of the marginal serial-dependence pattern into a perceptual component (green DoG curve, fitted to the partialled GLMM binned residuals) and a decisional component (purple sigmoid, fitted to the difference between the marginal and the perceptual binned residuals). Their algebraic sum is shown as the black dashed curve. The empirical marginal binned coefficients (data points) are read directly from the marginal GLMM and trace the predicted sum. Bottom row (panels c, d): empirical marginal DoG (solid coloured curves) and the marginal DoG fit to simulated data from the joint model (dashed black curves). For body fat the dashed and solid curves overlap closely throughout the Δ range shown. In muscularity the simulated DoG matches the bandwidth of the empirical marginal but has a smaller amplitude, consistent with the very wide perceptual channel in that block. The apparent narrow “channel-like” tuning of the marginal pattern (σ ≈ 1.4–1.8 z-units) is an emergent property of summing opponent processes with different widths, not a literal measurement of channel bandwidth.

#### Empirical marginal DoG (Figure 4 c and d, solid line)

Converting binary 2AFC responses into a continuous serial-dependence measure follows a standard residual-binning procedure. For each block we fit a GLMM in which the current response is regressed on the current stimulus, a set of indicator variables for one-unit-wide bins of Δ (with the Δ ≈ 0 bin as reference), and a per-subject random intercept. The Δ-bin coefficients are the empirical marginal serial-dependence effect on the log-odds scale at each Δ value, holding the current stimulus constant and averaging across observers. A positive coefficient at Δ > 0 (previous higher than current) indicates an attractive bias; a negative coefficient at the same Δ indicates a repulsive bias. We then fit the DoG and a linear model to these binned coefficients by weighted least squares (weights = 1 / SE).

At the marginal level (no partialling), the empirical DoG was strongly preferred over a linear model in both blocks (body fat: χ²(1) = 23.6, *p* < 10⁻⁵, ΔAIC = −21.6; muscularity: χ²(1) = 52.9, *p* < 10⁻¹², ΔAIC = −50.9). In body fat, the marginal DoG fit returned amplitude A = +0.282 (positive, indicating attraction) and bandwidth σ = 1.44 z-units. In muscularity, A = +0.337 and σ = 1.76 z-units. Both blocks show the textbook narrow attractive DoG, with σ ≈ 1.4 – 1.8 z-units corresponding to apparent tuned-channel bandwidths on the order of 14–18 body-number units. As we now show, this apparent bandwidth is not a measurement of underlying channel width.

#### Perceptual component (green DoG curve in **Figure 4a-b**)

We re-ran the binned GLMM with the centred previous response (*response_t-1 – 0.5*) added as an additional fixed-effects predictor. The Δ-bin coefficients from this joint GLMM now reflect only the influence of the previous stimulus, with the decisional carryover statistically removed. We fit a DoG to the resulting binned coefficients; the fitted curve is the green DoG in Figure 4 a and b. In body fat, the fitted amplitude was A_perceptual_ = –0.265 (negative, indicating repulsion) and the fitted bandwidth was σ_perceptual_ = 5.32 z-units, corresponding to a channel half-width of approximately 53 body-number units. In muscularity, the perceptual residual was essentially linear in Δ within the observable range: the DoG bandwidth saturated at the upper edge of the search grid (σ_perceptual_ ≥ 20 z-units, equivalent to a channel half-width exceeding 200 body-number units), with amplitude A perceptual = –0.242. The perceptual component is therefore broad and repulsive in both blocks, with the muscularity perceptual channel sufficiently wide that no curvature is detectable across the Δ range tested here.

#### Decisional component (purple sigmoid curve in **Figure 4a-b**)

The joint model contains a decisional term, *βprevious_response · (response_t–1 – 0.5)*, which is a linear function of the centred previous response. The decisional contribution to the log-odds on a given trial is therefore simply *βprevious_response* times the centred previous response (which equals *+βprevious_response /2* when the previous response was ‘high’ (i.e. ‘fat’ or ‘muscular’) and – *βprevious_response /2* when it was ‘low’ (i.e. ‘skinny’ or ‘scrawny’). The decisional component is “linearly modulated” in the sense that this contribution is a straight-line function of the previous response in log-odds space. It is not itself a DoG.

To plot the decisional contribution as a function of Δ, so that it can be compared visually to the perceptual DoG, we use the fact that the previous response is sigmoidally related to the previous stimulus through the observer’s own psychometric function: the probability that the previous response was ‘high’ given a particular previous stimulus follows the same logistic that describes current-stimulus discrimination. We therefore fit a smooth anti-symmetric sigmoid (i.e. the function takes opposite-signed values at equal distances either side of zero) to the empirical decisional contribution at each Δ (computed as the marginal binned coefficient minus the perceptual binned coefficient):

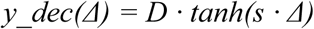

Here, the decisional contribution rises from *–D* at Δ ≪ 0 (where the previous response was usually ‘low’) through zero at Δ = 0 to *+D* at Δ ≫ 0 (where the previous response was usually ‘high’), with steepness *s*. For body fat judgments the fitted asymptote was *D* = +0.78 and the steepness, *s*, was = 0.69. For muscularity judgments, *D* = +1.10 and *s* = 0.46. The asymptotes correspond to *β_prev_resp* /2 (the log-odds shift between previous-‘high’ and previous-‘low’ conditions) and align closely with the joint-model *β_prev_resp* values (*β_prev_resp* /2 = +0.77 for body fat, +1.00 for muscularity).

#### Predicted sum (black dashed in Figure 4 a and b)

The predicted marginal serial dependence at each Δ is the algebraic sum of the perceptual and decisional contributions:

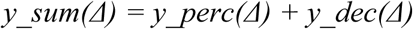

This sum is plotted as the black dashed line in Figure 4 a and b. By construction (since *y_dec* was fitted to marginal minus perceptual at each binned Δ), the sum closely tracks the empirical marginal binned coefficients (the red and blue dots), providing a direct visual confirmation of the two-process decomposition. The dashed curve lies within the error bars of the binned coefficients across the observable Δ range in both blocks.

#### Simulation-based recovery of the empirical marginal DoG (Figure 4 c and d, dashed line)

As an independent quantitative test of the decomposition, we simulated synthetic 2AFC responses from the fitted joint model and refitted the marginal DoG to the simulated data. For each actual trial in the dataset, we computed a predicted log-odds value by summing four contributions: a baseline calibrated to the empirical response rate, the current-stimulus contribution (*β_stim* times the trial’s current stimulus, where *β_stim* is the GLMM-fitted current-stimulus slope), the perceptual contribution (the fitted perceptual DoG evaluated at the trial’s Δ), and the decisional contribution (*β_prev_resp* times the trial’s centred previous response). We then converted the log-odds to a probability via the logistic function, and drew a simulated current response from a Bernoulli (coin-flip) trial with that probability. This yielded one simulated dataset of the same size and structure as the actual data. We then refit the binned marginal GLMM and the marginal DoG to the simulated dataset, exactly as we had for the empirical data.

In body fat, the simulated marginal DoG amplitude was *A* = +0.232 (empirical *A* = +0.282, difference = –0.050) and the simulated bandwidth was σ = 1.54 (empirical σ = 1.44, difference = +0.10). The simulation reproduces both the amplitude and the bandwidth of the empirical marginal to within sampling noise. In muscularity, the recovery was partial: the simulated DoG matched the empirical bandwidth well (σ_simulated_ = 1.89 vs σ_empirical_ = 1.76, difference = +0.13) but recovered only about 60% of the empirical amplitude (A_simulated_ = +0.199 vs A_empirical_ = +0.337, difference = –0.138). That the muscularity simulation does not fully reproduce the empirical amplitude likely reflects the very wide perceptual channel for that dimension (σ_perceptual_ saturating at the upper bound of the fit), which makes it harder to disentangle perceptual repulsion from decisional attraction precisely than in body fat. That said, the qualitative two-process decomposition holds in both blocks: the simulated marginal DoG closely matches the empirical bandwidth, and matches the empirical amplitude in both sign and order of magnitude.

The narrow DoG tuning of the marginal pattern (σ ≈ 1.5–1.8 z-units, or 15–18 body-number units) is therefore not a measurement of channel bandwidth: the underlying perceptual channels are at least three to four times broader in body fat (σ_perceptual_ ≈ 5.3 z-units, or about 53 body-number units), and broader still in muscularity (σ_perceptual_ large enough that the perceptual DoG is essentially linear within the observable Δ range). The apparent narrowness of the marginal pattern arises because a broad repulsive perceptual DoG and a broad attractive decisional sigmoid sum to a curve whose peak is sharper than either component on its own. That a simulation parameterised entirely by the joint-model coefficients reproduces the bandwidth of the empirical marginal DoG in both blocks (and the amplitude in body fat) is evidence that the two-process decomposition captures the bulk of the marginal pattern. No additional mechanism is required to explain the classic narrow attractive DoG. The apparent narrow tuning is an emergent property of summing opponent processes with very different widths.

### The decisional component is genuinely response-driven

To verify that the decisional component is genuinely driven by the previous response (rather than a hidden stimulus-driven effect aliased by the strong stimulus-response correlation), we ran two converging diagnostic tests (Tests 1 & 2), followed by a cell-level visualisation of the result (Test 3), using a clear-stimulus subset (body fat: body stimuli numbers < 50 or > 80, muscularity: stimuli numbers < 30 or > 60), within which each previous trial can be unambiguously classified as either correct or incorrect.

#### Test 1: Error perseveration

If the decisional component is genuinely response-driven, observers should carry forward their wrong responses on incorrect-previous trials, which would reduce current-trial accuracy after errors. We confirmed this prediction. In the body fat block, mean current accuracy was 0.973 after correct previous trials and 0.937 after incorrect previous trials; the difference (0.036) was significant in a per-subject paired test (t(57) = 2.52, *p* = .015, n = 58 subjects with both cell types). The effect was substantially larger in muscularity: 0.944 vs. 0.863, mean difference 0.081, paired t(70) = 4.63, *p* = 1.6 × 10⁻⁵, n = 71. Error perseveration is a textbook signature of response-level carryover and is mechanistically inconsistent with a purely perceptual account.

#### Test 2: Response repetition above chance after errors

Within the incorrect-previous subset, we computed the proportion of trials on which the current response repeated the previous (wrong) response, per subject. If the decisional component is genuinely response-driven, this proportion should exceed chance (0.5). In muscularity, the proportion was 0.708 (per-subject mean), significantly above 0.5: one-sample t(71) = 9.08, *p* = 1.7 × 10⁻¹³. In body fat, the proportion (0.514) did not differ significantly from chance (t(68) = 0.41, *p* = .69), consistent with the smaller decisional carryover coefficient in that block (βresponse = +1.53 in body fat vs +2.00 in muscularity). The response-repetition diagnostic therefore confirms the decisional account specifically in muscularity, where the decisional component is largest; the within-block ordering of effect sizes is internally consistent with the GLMM coefficients.

#### Test 3: Error trials - joint-model dissociation at the cell level

A more direct way to see the perceptual–decisional dissociation is through error trials. On most trials the previous stimulus and the previous response agree (a correct response), so the perceptual and decisional pulls reinforce each other and cannot be distinguished. The exceptions are the error trials, when the previous stimulus was ‘low’ but the observer responded ‘high’ (or vice versa), where the two signals point in opposite directions. If the joint model is right, the current judgment on these trials should follow the previous response, not the previous stimulus.

Figure 5 shows P(response = ‘high’) on the current trial at ambiguous current stimulus (near PSE), broken down by all four combinations of previous-stimulus category (low/high) and previous-response category (‘low’/’high’), within the stimulus subset. Subject counts above each bar reflect the inclusion criterion: a subject is included in a cell only if they produced at least one trial in that cell within the clear-stimulus subset. The correct-response cells therefore include nearly all subjects (73 in body fat, 72 in muscularity), while the rarer error cells include fewer (28 to 58, varying by cell and block). The asymmetry between the two error cells reflects a small response bias toward the ‘high’ response category (‘fat’/’muscular), more pronounced in body fat. In the error cells (light bars), the bias on the current trial follows the previous response: after a ‘low’ previous stimulus that received a ‘high’ response, P(high) on the current trial was 0.71 in body fat and 0.72 in muscularity; after a ‘high’ previous stimulus that received a ‘low’ response, P(‘high’) dropped to 0.43 and 0.26 respectively, a 28-to 46-percentage-point swing within the error cells alone. The correct-response cells (dark bars) show smaller, more balanced biases, because on these trials the perceptual and decisional pulls partially cancel each other. Fitting a categorical GLMM to these data confirms the dissociation statistically: the previous-stimulus and previous-response effects are opposite in sign and similar in magnitude in both blocks (body fat: –1.08 and +1.09 log-odds, both *p*-values < 10⁻¹¹; muscularity: –0.84 and +1.21, both *p*-values < 10⁻¹³).

**Figure 5.**
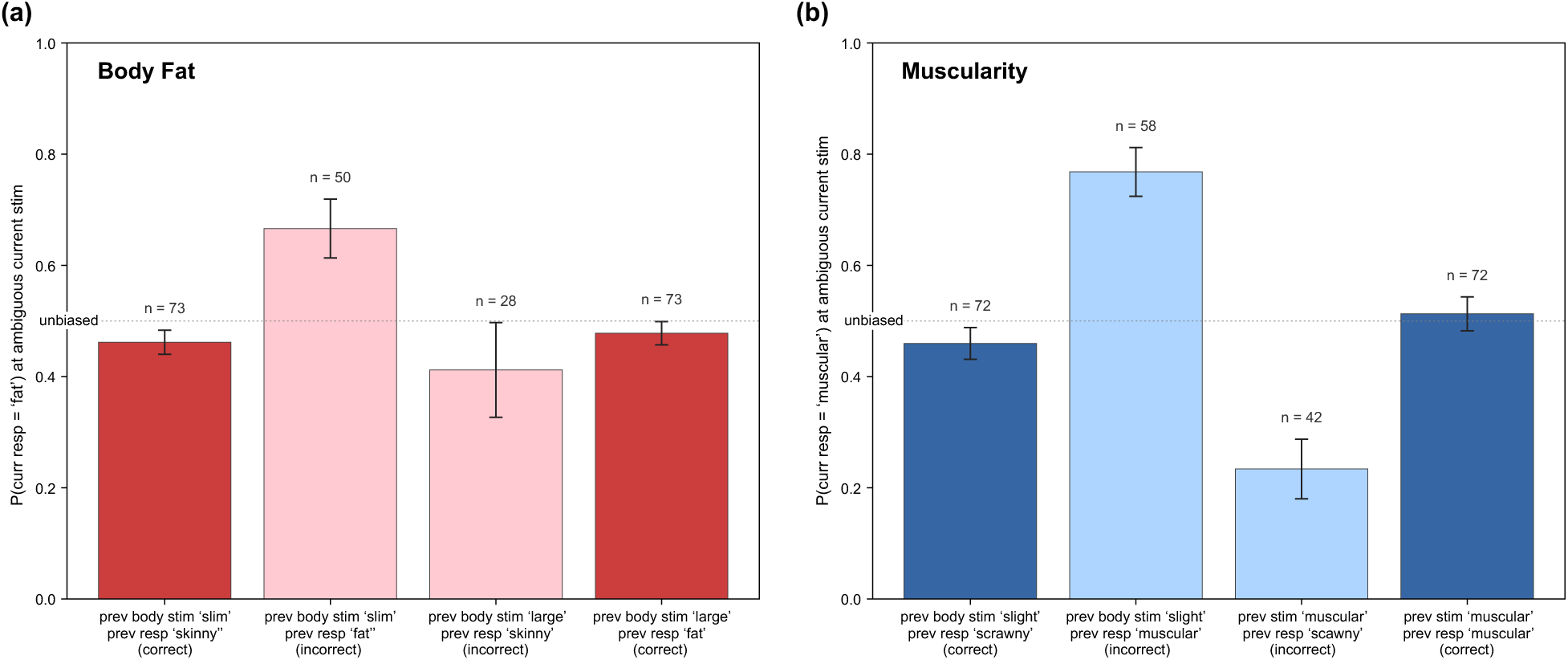
Cell-level visualisation of the perceptual–decisional dissociation through error trials. Per-subject P(response = ‘high’) on the current trial at ambiguous current stimulus (near PSE), broken down by the four combinations of previous-stimulus category (low/high) and previous-response category (‘low’/’high’), within the clear-stimulus subset. Light bars are error trials (previous stimulus and previous response mismatched) and dark bars are correct trials. The error cells show the largest biases, in opposite directions, with the current judgment following the previous response rather than the previous stimulus, which is the natural-experiment signature of the two-process decomposition. Subject counts (*n*) above each bar vary because a participant is included in a cell only if they produced at least one trial in that cell; the correct-response cells include nearly all participants, while the error cells include only the subset who made that particular error.

This analysis does not introduce a new inferential test: the same information is present in the joint GLMM (previous section), and Diagnostic 1 above already documented the same post-error phenomenon as a reduction in accuracy rather than as a directional bias. Its purpose is to make the dissociation directly visible: on error trials the perceptual and decisional pulls point in opposite directions, and the current judgment follows the response, not the stimulus.

### Long-range temporal persistence with dimensional differences

The joint model so far estimates carryover from the immediately preceding trial (*t*-1). A natural follow-up question is how long the perceptual and decisional traces persist and do the two components decay at the same rate?

To answer this, we re-fit the joint model at each lag from k = 1 to k = 30, replacing the previous-trial stimulus and response with the stimulus and response from k trials earlier. The model is otherwise identical to the lag-1 joint model used throughout. Each fit yields two coefficients: a perceptual coefficient (the lag-k stimulus effect, repulsive when negative) and a decisional coefficient (the lag-k response effect, attractive when positive). Figure 6 plots these coefficients as a function of lag for both blocks. In words, each dot is the strength of carryover from a single previous trial at the corresponding lag, holding the current stimulus and subject-level baselines constant.

**Figure 6.**
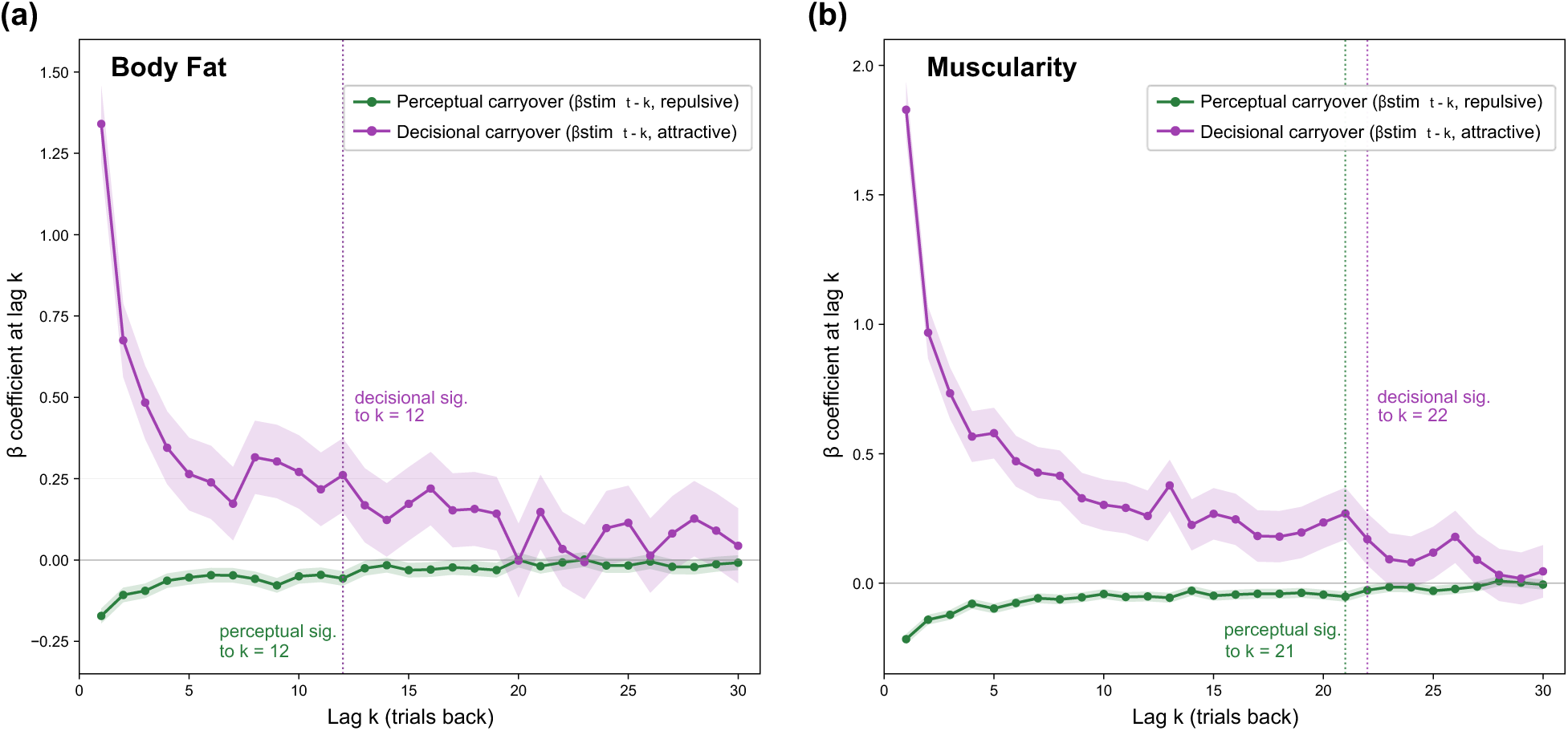
Long-range decay of the perceptual and decisional carryover components. Joint-model coefficients are plotted as a function of lag k (1 to 30 trials back) for body fat (left panel) and muscularity (right panel). Green: perceptual residual (the lag-k stimulus coefficient, negative because perceptual carryover is repulsive). Purple: decisional carryover (the lag-k response coefficient, positive because decisional carryover is attractive). Shaded bands are 95% confidence intervals. Vertical dotted lines mark the largest lag at which each component remained statistically significant under a Bonferroni-corrected threshold of α/30 = 0.0017 within block, tolerating a one non-significant intermediate lag. Body fat: perceptual residual significant to k = 12, decisional to k = 12. Muscularity: perceptual residual significant to k = 21, decisional to k = 22.

Three features stand out. First, both components decay monotonically with lag: trials further in the past have progressively less influence on the current response, with a sharp initial drop from lag 1 to lag 2 and a slower decline thereafter. Second, the decay is much slower in muscularity than in body fat. In body fat, the perceptual residual remains statistically significant out to lag k = 12, and the decisional carryover also out to lag k = 12 (Bonferroni-corrected within block at α/30 = 0.0017, tolerating a single non-significant intermediate lag). In muscularity, the corresponding limits are lag k = 21 (perceptual) and lag k = 22 (decisional). Both components, therefore persist roughly 75 to 85 percent longer in muscularity than in body fat. Third, the qualitative shape of the decay is the same in both blocks, a smooth monotonic decline with no abrupt drops or oscillations. This argues against a dimension-specific reset or boundary mechanism: the same processes appear to operate in both blocks, simply on different timescales.

The slower decay in muscularity is consistent with the channel-encoding conclusion above: broader, less specific perceptual channels would be expected to recover more slowly than narrowly tuned ones.

### Tuned-channel architecture for body fat, but not muscularity

Having isolated the perceptual component of serial dependence from the decisional component, we can now ask a more precise question about the structure of that perceptual effect. Does repulsion vary with the feature distance between inducer and target in a way that reflects discrete, bandwidth-limited channels, or does it simply grow (or shrink) smoothly with distance, with no hint of channelled organisation?

To answer this, we fitted two competing models to the perceptual residual as a function of Δ (the signed feature difference between the previous trial’s stimulus and the current one). The perceptual residual here is the trial-by-trial log-odds of a “higher” response after partialling out the previous response, so it captures the purely stimulus-driven carryover: the tendency for the current percept to be pushed away from where the preceding stimulus sat. This is what the y-axis in Figures 7a and 7b indexes. A positive value means the observer is more likely to respond “higher” when the inducer was lower than the target (repulsion away from the inducer), and a negative value means the reverse. In short, it measures how strongly, and in which direction, the ghost of the last trial biases the current percept. The question is whether this repulsive influence is shaped (rising and then falling as the inducer moves further away) or simply linear.

**Figure 7.**
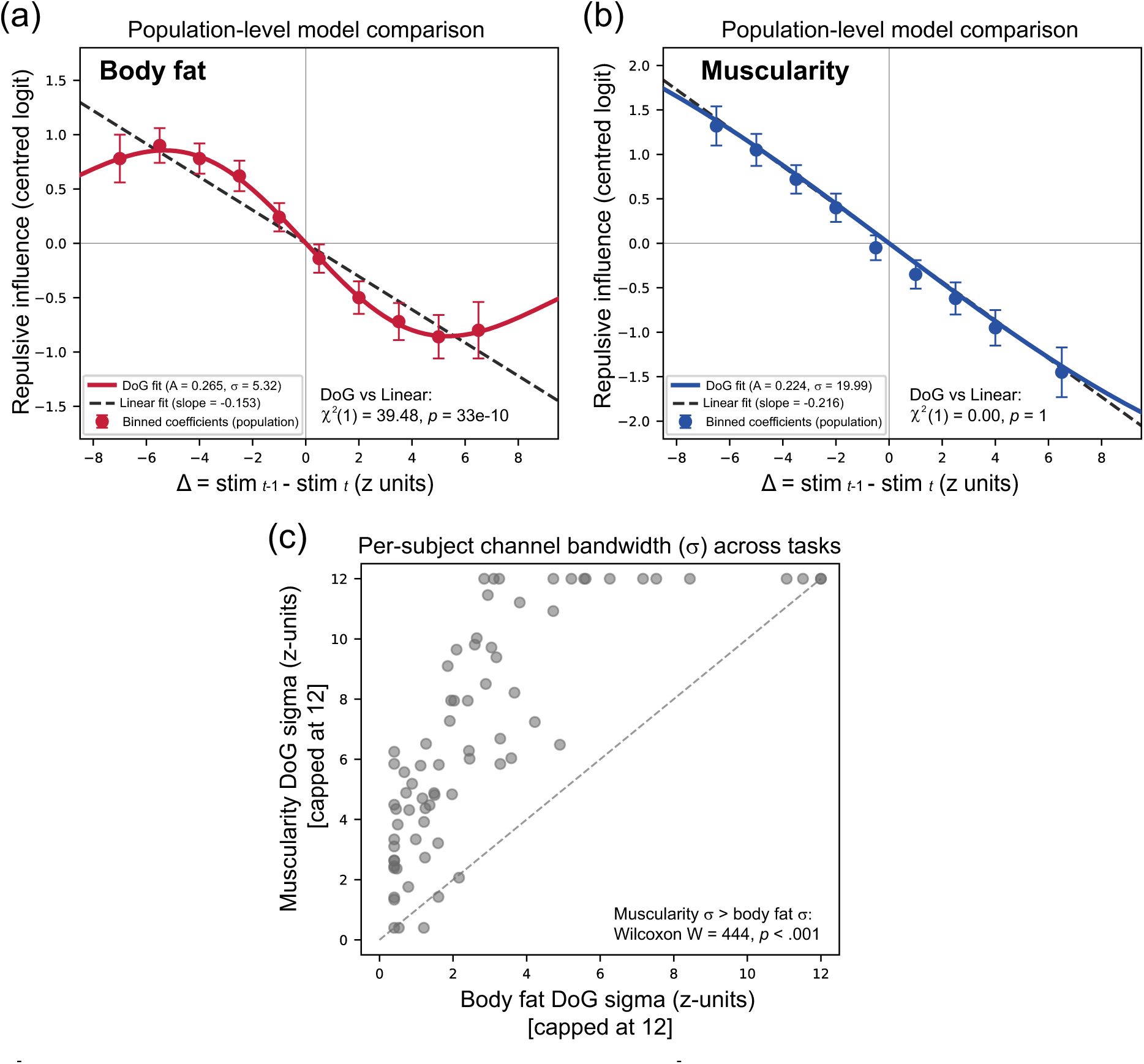
Channel-tuning architecture: DoG versus linear model comparison. Population-level model comparison for body fat (a) and muscularity (b). Coloured symbols are binned coefficients from the joint model (previous response partialled out). Solid coloured lines are DoG fits and the dashed lines are linear fits. (c) Per-subject paired comparison of fitted bandwidth σ, showing that muscularity channels are systematically broader than body fat channels.

The two models are a linear function (repulsion proportional to Δ, with no channel structure) and a Difference-of-Gaussians (DoG) function, which produces exactly the peaked-then-declining shape expected when a tuned channel is selectively adapted by the inducer. We compared them using likelihood-ratio tests, pooling trials across all 74 participants within each block, and fixing the previous-response term so that the decisional component remained controlled throughout.

The two body-composition dimensions produced strikingly different results. For body fat, the DoG model fitted substantially better than the linear model: χ²(1) = 39.48, p = 3.3 × 10⁻¹⁰, ΔAIC = −37.5.

The estimated channel bandwidth was σ = 5.32 z-units (roughly 53 body_num units), and the adaptation amplitude was A = +0.265 (positive A indicates repulsion under our sign convention). The shape of this effect, visible in Figure 7a, is diagnostic: repulsion peaks at intermediate inducer-target separations and falls off at large ones, exactly as a tuned-channel model predicts. For muscularity, the DoG model provided no improvement whatsoever: χ²(1) ≈ 0, ΔAIC = +2.1 (the degrees of freedom penalty actually favours the simpler linear model). The fitting pushed σ to its upper boundary (σ = 20 z-units), at which point the DoG function becomes indistinguishable from a straight line across the range of Δ values we observed. The muscularity data are therefore fully consistent with a graded, linear repulsive aftereffect, with no detectable channel-tuning structure (Figure 7b).

The same pattern is evident within individuals. Fitting the DoG model separately for each participant and each block, we recovered per-subject σ estimates (capped at 12 z-units where the bandwidth was unidentifiable given the trial count). Mean σ was 6.16 z-units for body fat and 9.03 z-units for muscularity, and a paired Wilcoxon test confirmed that this within-person difference is systematic: W = 444, *p* = .0001, n = 74. Muscularity channels are reliably broader, or more precisely, less narrowly tuned, than body fat channels at the individual level (Figure 7c).

This comparison is worth making because it links the functional form of serial dependence to the representational architecture inferred from classical psychophysics. Narrow tuning in body fat implies that the perceptual system maintains distinct channels for body fat ranges, and that adapting one channel drives perception away from its preferred value. The absence of equivalent tuning in muscularity implies that muscularity is encoded without the same degree of channel discretisation, at least at the resolution probed by our stimulus range. The dimensional dissociation is therefore not just a quantitative difference in effect size: it is a qualitative difference in the kind of representation that generates serial dependence.

A further analysis strengthens the channel-based interpretation for body fat. When observers are split at the median JND within each block and the distance-tuning function is re-estimated separately for each subgroup, a predicted pattern emerges (Figure 8a, 8b). In the body fat block, low-JND observers (better discriminators, who by the channel hypothesis should have narrower bandwidths) show a more peaked DoG, with the repulsive peak concentrated around Δ ≈ 4 z-units. High-JND observers show an effectively linear function, consistent with channels too broad for the DoG signature to be visible within the observable Δ range. This contrast is absent in muscularity, where both JND subgroups produce essentially linear functions (Figure 8b). The 3-way interaction (task × JND subgroup × DoG shape) is highly significant, χ²(8) = 569.4, *p* < 10⁻¹¹⁵, with the JND-related amplitude difference 2.3× larger in body fat than in muscularity.

**Figure 8.**
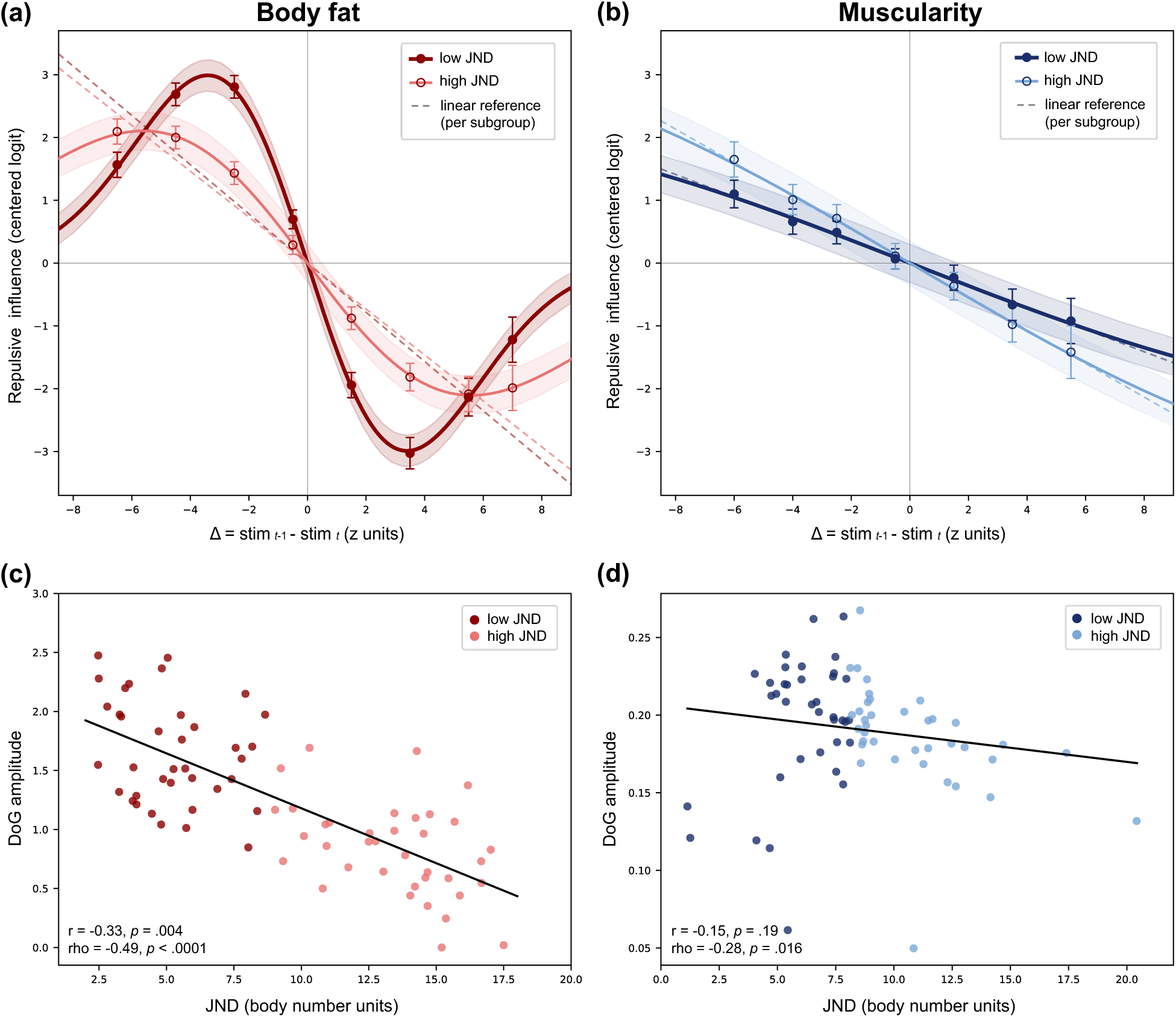
Feature distance-tuning split by JND group, and DoG amplitude versus individual sensitivity. Top row: distance-tuning of the perceptual residual for body fat (a) and muscularity (b), with observers split at the median JND within each block. Dark curves and filled symbols represent low-JND observers, light curves and open symbols high-JND observers. Solid lines are DoG fits, dashed lines are the matched linear reference for each subgroup, shaded bands are 95% confidence intervals, and points are binned Δ coefficients. Bottom row: per-subject DoG amplitude A against individual JND for body fat (c) and muscularity (d).

### Individual differences in the body-fat perceptual channel reflect adaptation of amplitude, not bandwidth

Before turning to individual differences, we first established that the body-fat channel signature is genuinely a within-observer property and not a by-product of averaging across observers. This matters because a curved population-level tuning function can in principle arise artificially if observers differ in the slope of an underlying linear effect, so that pooling their data manufactures apparent curvature where none exists within any individual. We therefore tested for the DoG curvature within each observer directly. For each participant, we partialled the linear trend out of the DoG-shaped predictor and estimated the coefficient on the orthogonal non-linear component. These per-subject coefficients were then submitted to a one-sample test against zero. The residual curvature was reliably repulsive, with a mean coefficient significantly below zero (t(73) = 3.87, *p*= 2.3 × 10⁻⁴; Wilcoxon *p* = 3.4 × 10⁻⁴), with 49 of 74 observers (66%) exhibiting the DoG curvature in the predicted direction. Because the DoG component is orthogonal to each observer’s own linear fit, it cannot be a pooling artefact. The perceptual component’s DoG shape arises within individual observers rather than just across them.

Having established that the per-observer channel signature is real, we asked which parameter of the tuning function accounts for individual differences in discrimination ability. Two parameters of the DoG could in principle vary across observers: the amplitude A (the magnitude of adaptation) or the bandwidth σ (the channel width). We fitted per-subject DoG functions to the body-fat perceptual residual and tested both parameters against per-subject JND (Figure 8c). Adaptation amplitude (A) was strongly and inversely related to JND in the body fat task: r = −0.33 (*p* = .004) and ρ = −0.49 (*p* = 1.1 × 10⁻⁵), n = 74. Better discriminators (lower JND) showed roughly twice the adaptation amplitude of poorer discriminators. Channel bandwidth σ, by contrast, showed no relationship with JND: r = −0.17 (*p* = .22) within the subset of observers whose σ was identifiable in body fat. The population-level analysis tells the same story. When low- and high-JND subgroups are fitted separately (Figure 8a), the amplitude differs by about a factor of two (low-JND A = 0.59 versus high-JND A = 0.30), whereas the bandwidth differs by only about 30% (low-JND σ = 4.38 versus high-JND σ = 6.15).

In muscularity, the corresponding relationships were weak (Figure 8d) and did not support a channel-tuning account. Amplitude against JND was not significant by Pearson correlation (r = −0.15, *p* = .19), with only a weak negative trend by the rank-based test (ρ = −0.28, *p* = .016). σ against JND was likewise null. As noted earlier, the muscularity amplitude estimate is difficult to measure because the bandwidth runs to its upper bound in most observers. We therefore do not interpret this weak rank-order trend as evidence for a channel structure that the population-level analysis has already failed to detect.

### Convergent evidence for tuned-channel encoding of body fat, but not muscularity

Several converging analyses support the conclusion that body fat, but not muscularity, is mediated by tuned perceptual channels:

1. Population-level DoG versus linear model comparison: the DoG is strongly preferred over a linear fit for body fat (χ²(1) = 39.48, *p* < 10⁻⁹), with no improvement over linear for muscularity (*p* = 1) (Figure 7).
2. Per-subject paired bandwidth (σ) comparison: muscularity bandwidths are significantly broader than body fat bandwidths (Wilcoxon W = 444, *p* = .0001, n = 74) (Figure 7c).
3. JND-moderated distance-tuning shape: the 3-way task × JND × shape interaction is highly significant (LRT χ²(8) = 569.4, *p* < 10⁻¹¹⁵), with the JND effect 2.3× larger in body fat. Low-JND body-fat observers show a narrower DoG, whereas muscularity is linear in both JND subgroups (Figure 8a,b).
4. Adaptation amplitude versus JND in body fat: per-subject DoG amplitude correlates inversely with JND (r = −0.33, *p* = .004; ρ = −0.49, *p* = 1.1 × 10⁻⁵), consistent with channel adaptation magnitude varying with discrimination ability; the relationship is null or uninterpretable in muscularity (Figure 8c).

### Effects of response/viewing time

Because the task was response-terminated, the time for which each body remained visible varied from trial to trial, and the response time on a trial is also its stimulus exposure duration. The median viewing time was 581 ms in the body fat task and 619 ms in the muscularity task, with both distributions right-skewed (Figure 9a). The two distributions overlapped almost completely, so comparisons between the dimensions are not confounded by differences in viewing time.

**Figure 9.**
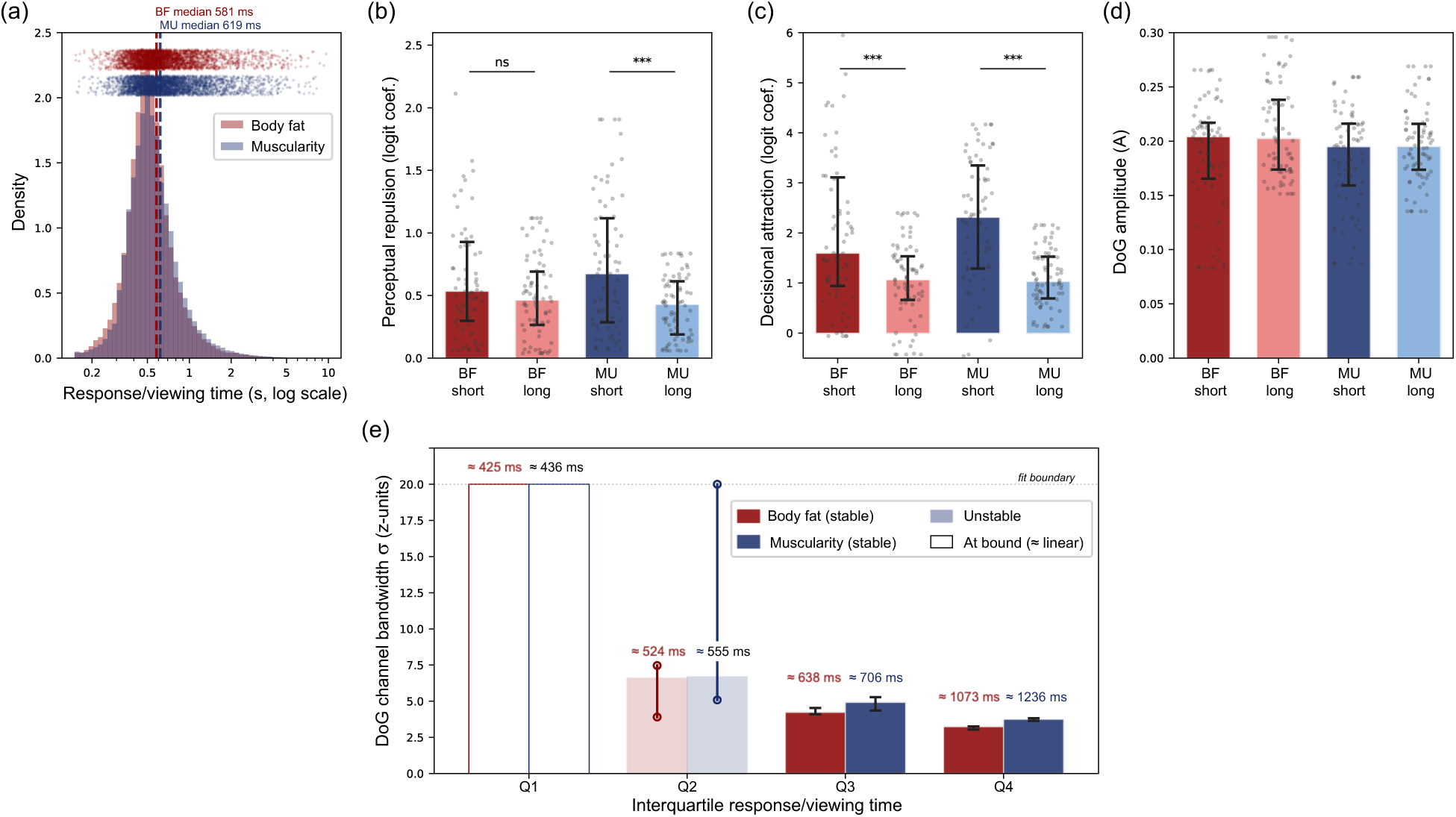
Response time as viewing time: modulation of serial-dependence components. Because the task was response-terminated, a trial’s response time equals the time the body was on screen. (a) Distributions of response (viewing) time for the body fat (red) and muscularity (blue) tasks, on a log scale. Dashed lines indicate the medians (581 and 619 ms). Raw trials distributions are shown above. The two distributions overlap almost completely. (b, c) Perceptual repulsion and decisional attraction, estimated separately for each observer’s short-viewing and long-viewing trials (split at their own median response time within each task). Bars show the median across observers and error bars the interquartile range; grey points are individual observers. Brackets show Mann-Whitney comparisons of short versus long (ns, not significant; *** *p* < .001). Decisional attraction is larger on short-viewing trials in both tasks; perceptual repulsion is larger on short-viewing trials for muscularity but not body fat. (d) Amplitude of the perceptual DoG for each task and viewing-time group (median ± IQR). (e) Bandwidth (σ) of the perceptual DoG estimated within each observer’s response-time quartile, with the median viewing time of each quartile shown above it (red = body fat, blue = muscularity). Solid bars are stable estimated bandwidths, open bars at the upper limit (“at bound”) indicate quartiles in which the fit ran to the edge of the search space and the function is effectively linear; faint bars with whiskers (“unstable”) indicate quartiles in which the estimate varied. Bandwidth narrows as response/viewing time increases, from unidentifiable in the fastest quartile (∼430 ms) to roughly 4 to 5 z-units in the two slowest quartiles (∼670 and ∼1150 ms), in both dimensions.

We first confirmed that the perceptual and decisional components are not artefacts of viewing time. Adding the log of the current and previous response times as covariates to the joint model left both coefficients essentially unchanged (perceptual coefficient altered by less than 0.3%, decisional coefficient by less than 1.5% in both tasks), so neither component is driven by how long the stimulus was seen.

We then asked whether viewing time modulates the size of each component, splitting each observer’s trials at their own median response time (Figure 9b, c). Decisional attraction was larger on short-viewing than long-viewing trials in both tasks (Mann-Whitney, both *p*-values < .001). Perceptual repulsion showed the same direction but reached significance only for muscularity (p < .001), not body fat (*p* = .09). The larger decisional carryover on fast trials is consistent with the well-documented tendency for quick responses to repeat the previous choice.

The amplitude of the perceptual DoG was unaffected by viewing time, and was comparable across both tasks and both viewing-time groups (A ≈ 0.20 throughout; Figure 9d). The channel bandwidth, in contrast, depended strongly on viewing time. Estimating the DoG bandwidth within each observer’s response-time quartile (Figure 9e) showed that it narrowed as viewing time lengthened: in the fastest quartile (median viewing time ≈ 430 ms) the bandwidth was unidentifiable, running to the upper bound of the search in both tasks, whereas in the two slowest quartiles (≈ 670 and ≈ 1150 ms) it settled to well-constrained values of roughly 4 to 5 z-units. This trend was present in both dimensions, with body fat slightly more narrowly tuned than muscularity at each quartile in which bandwidth could be estimated. Notably, muscularity, whose perceptual residual is linear in the aggregated data, yielded a clearly curved, channel-like residual under the longer viewing times.

Two things follow from this. First, the perceptual tuning width is estimable only when viewing is long enough; in the fastest trials the bandwidth cannot be resolved, in part because the perceptual residual on those trials is left after removing a larger and noisier decisional component. Second, because response time and exposure duration are the same quantity in this design, we cannot say whether the narrowing of bandwidth with viewing time reflects exposure duration, the smaller decisional component at longer response times, or both. We return to this in the Discussion.

## Discussion

We applied a trial-level partialling framework to a 2AFC body-discrimination dataset in order to separate the perceptual and decisional contributions to serial dependence, and to ask whether each of two body-composition dimensions, body fat and muscularity, carries the signature of tuned-channel encoding. The perceptual/decisional decomposition of serial dependence (Pascucci et al., 2019) generalises to body perception, and the weak attractive bias seen in the raw data proves to be the near-cancellation of two much larger opposing processes. Body fat shows the channel signature predicted by adaptation accounts, but muscularity does not (in the aggregate data), a dimensional dissociation we did not anticipate. And where the channel signature is present, individual differences in discrimination ability are carried by the strength of adaptation rather than the width of the channels. We discuss these findings in sequence, then consider the methodological advance that made them visible, their possible neural basis, and their broader implications for everyday vision and for future work.

### Perceptual and decisional components of serial dependence generalise to body perception

At the marginal level, serial dependence in our data was weak (albeit highly statistically reliable) and slightly attractive: the previous stimulus exerted a small positive influence on the current response, with coefficients of about +0.04 in body fat and +0.08 in muscularity. Taken at face value, this would suggest little carryover, and it would sit uneasily beside the substantial attractive biases previously reported for body size (Alexi et al., 2018; Alexi et al., 2019). Our trial-level partialling shows that this apparent near-absence is misleading. When the previous response was added to the model, the previous-stimulus coefficient reversed sign in both dimensions, to roughly−0.22 in body fat and −0.25 in muscularity, revealing a strong perceptual repulsion: once decisional carryover is held constant, the current body is pushed away from the preceding one. At the same time, a large positive previous-response coefficient emerged, about +1.5 in body fat and +2.0 in muscularity, indicating a strong tendency to repeat the previous response. The flat marginal pattern is therefore the sum of a large repulsive perceptual component and a large attractive decisional component that conceal one another.

This opposite-signed structure is now an established and general property of serial dependence. The dissociation was first shown for orientation by (Fritsche et al., 2017) and formalised by (Pascucci et al., 2019), and across the broader literature it has been documented in working memory as well as orientation, such that the reported judgment is commonly drawn toward the prior response, a tendency termed decisional carryover, while the underlying perceptual representation is repelled (reviewed in (Manassi et al., 2023); see also (Gallagher & Benton, 2024)). To our knowledge, this is the first demonstration that serial dependence in visual body perception comprises two components of opposite sign, since all prior body serial-dependence work has used reproduction tasks (the bodyline) and has not separated them (Alexi et al., 2018; Alexi et al., 2019; Turnbull et al., 2022). A set of converging diagnostics confirmed that the decisional component is genuinely response-driven rather than a stimulus effect in disguise (see decisional diagnostic tests in Results): accuracy fell after error trials, responses repeated above chance following errors, and on error trials, where the previous stimulus and the previous response point in opposite directions, the current judgment followed the response rather than the stimulus.

The channel-based account predicts that both body fat and muscularity, having been shown to adapt independently and dimension-specifically under prolonged adaptation (Brooks, Keen, et al., 2020) (Sturman et al., 2017), should carry the DoG signature in their perceptual residuals. The DoG-shaped feature tuning of serial dependence, and the interpretation of its bandwidth as a tuning estimate, are themselves well established and have been characterised across orientation, face, and numerosity perception (Fischer & Whitney, 2014); reviewed in (Manassi et al., 2023). Only body fat met this prediction in our aggregated data set. For body fat the DoG fitted the perceptual residual substantially better than a straight line, with repulsion that peaked at intermediate inducer-target distances (bandwidth σ ≈ 5.3 z-units, about 53 body-number units) and declined at larger ones, the expected fingerprint of a tuned channel. For muscularity the perceptual repulsion was robust but monotonic and linear across the tested range, with no detectable peak and a bandwidth that ran to the upper edge of the search space, the limiting case in which a DoG is indistinguishable from a line.

The dissociation held within individuals as well as at the group level. Per-subject bandwidths were reliably narrower for body fat than for muscularity, and the body-fat curvature was reliably present within single observers once each observer’s own linear trend was removed, with about two thirds of observers showing curvature in the repulsive direction. It is therefore not a by-product of averaging across heterogeneous people. The two dimensions also differed in time course, with both carryover components persisting markedly longer for muscularity than for body fat. This is consistent with the channel interpretation: broader, less specific channels would be expected to renormalise more slowly.

This dissociation, however, is not fixed across viewing conditions. Because the task was response-terminated, the time for which each body remained visible varied from trial to trial, and we used this to ask whether the perceptual channel estimate depends on how long the current body is seen. Sorting each observer’s trials into within-subject response-time quartiles and re-estimating the perceptual residual’s DoG bandwidth within each (Figure 9e) revealed a systematic trend: the bandwidth narrowed as viewing time lengthened. In the fastest quartile (median viewing time ≈ 430 ms) the bandwidth was the bandwidth was unidentifiable, reaching the upper bound of the search space. In the two slowest quartiles (≈ 670 and ≈ 1150 ms) it settled to well-constrained values of roughly 4 to 5 z-units. This pattern was present in both dimensions, with body fat slightly more sharply tuned than muscularity at every quartile in which bandwidth could be estimated. The DoG amplitude, in contrast, was comparable across all quartiles and both dimensions (Figure 9d), so what changed with viewing time was the apparent width of the channel rather than the strength of the adaptation. One point is worth noting, and we develop it cautiously below: at the longer viewing times the two dimensions are similarly tuned, so the difference between them in the aggregate may reflect how readily tuning can be resolved under brief viewing rather than a stable difference in channel width.

This is the point of contact with the work of Brooks and colleagues, who reported tuned, dimension-specific channels for both fat and muscularity using prolonged adaptation (Brooks, Keen, et al., 2020; Sturman et al., 2017). In our aggregated data the trial-by-trial method confirms tuned channels for body fat but not for muscularity. Taken alone this seems to conflict with their muscularity result. The contrast between the two dimensions is not one of the strength of the perceptual effect. The repulsion itself is essentially as large for muscularity as for body fat, with comparable DoG amplitudes (A ≈ 0.20 in both; Figure 9d). What differs is whether that repulsion is tuned: for body fat the perceptual residual is fit substantially better by a DoG than by a line, whereas for muscularity the residual is well described by a straight line, with the DoG bandwidth running to the upper bound of the search space, a range within which a DoG is indistinguishable from linear repulsion. The muscularity difference is therefore a failure to resolve tuning, not a ‘weaker’ effect. We are cautious about reading too much the response/viewing time analysis. The perceptual residual’s bandwidth was estimable only in slower trials and became unstable in faster ones (Figure 9e). Although muscularity’s residual is somewhat more stable under longer viewing, the bandwidth is precisely the parameter that is least stable here, and we do not treat the long-response/viewing time fits as firm evidence of a muscularity channel. The most we can say is that prolonged adaptation, which involves accumulated exposure over many seconds to minutes (in (Sturman et al., 2017), an initial 120 s of adaptation with top-ups between test trials), lies far beyond the longest single-trial exposures in our task, and that any tuning in muscularity that our method cannot resolve at these brief exposures might still be expressed under the much longer exposures of the adaptation paradigm. We therefore interpret the dimensional difference as one of the statistical *resolvability* of channel tuning under brief viewing, rather than as the categorical presence of tuned channels for body fat and their absence for muscularity, while acknowledging that our aggregated data establish a tuned channel only for body fat.

### Individual differences reflect adaptation strength, not channel width

For body fat, where a clear channel signature exists, we asked which property of the tuning function accounts for differences between observers in discrimination ability. Two are available from our analyses: the amplitude of the DoG, which indexes how strongly the channel adapts, and the bandwidth, which indexes how broadly it is tuned. Discrimination ability predicted amplitude, not bandwidth. Better discriminators, those with lower JNDs, showed roughly twice the adaptation amplitude of poorer discriminators, while bandwidth was unrelated to JND, and the same pattern appeared at the population level when observers were split by JND. This pattern suggests a gain-control mechanism, whereby observers differ in how strongly their body-fat channels adapt to recent input, rather than in how finely those channels are tuned. The same decomposition cannot be applied to muscularity. Its bandwidth is unresolvable in the aggregated data (Figures 7-8), so amplitude and bandwidth cannot be separated. This reflects the difficulty of resolving its tuning, not a weaker effect (Figure 9e).

The methodological contribution of this work concerns what can be recovered from a 2AFC categorisation task. Serial dependence has been measured in binary and forced-choice tasks before, notably in audiovisual timing and temporal-recalibration tasks (Cass et al., 2026; Van der Burg et al., 2013; Van der Burg & Goodbourn, 2015). Estimates of the *tuning width* of the perceptual component, its derivative-of-Gaussian (DoG) bandwidth, have come from a different kind of task: principally from reproduction and adjustment paradigms, in which the observer regenerates a graded stimulus value on each trial and the continuous response itself traces the tuning function (Fischer & Whitney, 2014; Fritsche et al., 2017; Pascucci et al., 2019) reviewed in (Manassi et al., 2023). Such paradigms are both time-consuming and intrinsically noisy, since each reproduction carries motor and memory error. What has not, to our knowledge, been done is to recover the DoG bandwidth of the perceptual component from binary categorisation data. Two features of the present design make this possible. First, by including both the previous stimulus and the previous response as predictors in a trial-level mixed-effects model, the partial coefficient on each isolates one source while holding the other constant, so the perceptual residual can be separated from the decisional carryover within the same binary responses. Second, because the stimulus dimension is sampled across many graded levels of each stimulus dimension rather than a single comparison against a fixed standard, the perceptual carryover can be measured as a function of the featural distance between the current and previous trial(s). It is this featural distance profile that makes the perceptual bandwidth estimable. What is new here is therefore neither the two-component structure nor DoG tuning as such, both of which are well-established and general properties of serial dependence, but the estimation of the perceptual tuning width from a rapid forced-choice categorisation task.

The approach is also fast. Each block of 480 trials took, on average, about fifteen to twenty minutes to complete. Consequently, a full tuning profile for one dimension can be obtained in roughly the time of a brief questionnaire. This combination of speed and the ability to separate perceptual from decisional contributions is what recommends the method for settings, including clinical ones, where lengthy reproduction sessions are impractical.

The decomposition also has an important theoretical implication. The classic attractive serial-dependence pattern follows a narrow DoG, and this narrowness has often been read as a direct measurement of channel width. Our results show that it is not. At the marginal level, before the decisional carryover is removed, the DoG was narrow in both dimensions, on the order of 15 to 18 body-number units. The partialled perceptual residual is far broader wherever its width can be resolved. For body fat its repulsive tuning has a bandwidth of about 53 body-number units, several times the marginal value. For muscularity the aggregated residual is linear, with no resolvable peak, but when trials are sorted by viewing time it too resolves into a curved DoG at the longer exposures (Figure 9e), of broadly comparable, large bandwidth. In neither dimension, then, does the underlying perceptual channel approach the narrowness of the marginal pattern. A simulation built entirely from the fitted joint-model components reproduced the narrow marginal DoG, confirming that the marginal pattern is the algebraic sum of the broad (or, for muscularity in the aggregate, linear) repulsive perceptual component and the attractive decisional component. The apparent narrowness of the marginal DoG is therefore an emergent property of summing two opposing processes, not a property of any underlying channel. This provides a trial-level, quantitative confirmation of the sum-of-components account proposed by (Pascucci et al., 2019), and it shows directly why the marginal pattern misrepresents channel architecture, and why the partialled perceptual residual, not the marginal DoG, is the appropriate measure for testing channel-based encoding.

### Putative neural mechanisms

Our behavioural decomposition can be related to a growing body of neural work. We frame these links as hypotheses to be tested, not as claims our data establish. For the perceptual component, the most direct candidate substrate is body-selective occipitotemporal cortex, and specifically the fusiform body area (FBA). Using fMRI adaptation with thin and fat body images, Hummel et al. (2013) found that the FBA, together with the right middle occipital gyrus, showed neural adaptation to specific body shapes, whereas the extrastriate body area (EBA) did not, and that this adaptation was accompanied by a behavioural body-shape aftereffect. This makes the FBA a strong candidate locus for the repulsive perceptual component we isolate, since adaptation in body-shape-tuned populations is exactly the mechanism a channel account requires. If body fat and muscularity differ in whether they are channel-coded, as our results suggest, one possibility, testable with fMRI adaptation, is that fat and muscularity adapt to differing extents within these regions.

The relationship between the two components also aligns with the emerging neural picture of serial dependence as a two-stage process. Imaging studies have found that the representation of the current stimulus in early visual cortex is repelled from the recent past, consistent with adaptation, even while the behavioural report is attracted to it (Hajonides et al., 2023; Sheehan & Serences, 2022). This is generally interpreted as a repulsive bias at sensory encoding that is later counteracted by an attractive bias at a post-perceptual decision stage, with the attractive component linked to frontoparietal and prefrontal cortex (reviewed in (Manassi et al., 2023). This is the same architecture our decomposition recovers behaviourally: a stimulus-driven repulsion attributable to adaptation, and a response-driven attraction attributable to a decisional process. The correspondence suggests that the perceptual residual we isolate may index adaptation in sensory (for bodies, body-selective) cortex, while the decisional carryover may index the decision-stage contribution. Confirming this would require running the decomposition alongside neural measurement in the same observers.

### Implications for vision under natural conditions

The adaptation that Brooks and colleagues characterised with prolonged exposure, the sustained viewing of an adapting body over seconds to minutes, turns out also to operate from a single preceding trial. Our results show that body-composition channels are renormalised by a single preceding trial, on which the body was visible for less than two seconds in most cases, not only by prolonged exposure. For everyday vision, this implies that body perception is continuously recalibrated, each body slightly reshaping the appearance of the next, at a timescale far briefer than prolonged adaptation suggests. The repulsive direction of the perceptual effect fits an efficient-coding interpretation, in which rapid recalibration reduces redundancy and keeps channels centred on the recent input distribution (Cicchini et al., 2018; Sheehan & Serences, 2022), rather than a stabilising one. Whether this rapid recalibration and the slow aftereffect of prolonged adaptation reflect a single mechanism operating across timescales, or two distinct processes, is an open question that our data raise but cannot settle, since we did not measure the two within the same observers. We take it up as a future direction below.

### Why do body fat and muscularity judgments differ in their apparent tuning?

The most parsimonious account of the difference is experiential. Tuned channels are thought to arise through exposure, a dimension acquiring narrowly tuned channels to the extent that it has been encountered varying densely and meaningfully over the lifespan. Body fat varies visibly across nearly everyone an observer meets and is salient from early life, exactly the kind of richly sampled dimension for which sharp channels would be expected to form. Muscularity is encountered across a narrower range, is often obscured by clothing, and becomes a focus of attention later and for fewer observers, which could leave its tuning weaker or harder to resolve. This is a hypothesis, not something our data demonstrate. Our data establish a difference in the aggregate, not its origin, and at longer viewing times the two dimensions are similarly tuned (Figure 9e), so the difference may lie more in the resolvability of tuning under brief viewing than in channel width itself. If exposure underlies these differences, the account makes several testable predictions. Observers with extensive experience attending to male muscularity (such as individuals involved in strength sports or bodybuilding, or potentially men attracted to men, whose visual attention to and evaluation of male bodies may differ systematically from heterosexual men) may exhibit more robust or more readily measurable muscularity tuning that the general population sampled here.

### Relation to prior work and the contribution of this study

We do not claim priority for serial dependence in body perception (Alexi et al., 2018), nor for the perceptual/decisional decomposition, (Fritsche et al., 2017) and, in its fullest form (Pascucci et al., 2019), nor for the DoG tuning function and its use as a bandwidth estimate, which are standard tools used since Fischer and Whitney (2014) and reviewed across domains by Manassi et al. (2023). Our contribution lies in the combination of these elements and in three specific results. First, we provide a within-subject test of channel-based encoding across two orthogonal body-composition dimensions, using the partialled perceptual residual rather than the contaminated marginal pattern as the measure, and we estimate the channel-tuning profile from a 2AFC categorisation task, which to our knowledge has not been done before, since previous derivations have required reproduction or estimation data. Second, we provide a trial-level, quantitative confirmation of the Pascucci et al. (2019) sum-of-components account, showing by simulation that the marginal DoG is closely reproduced by the fitted components. Third, we provide the first characterisation of serial dependence for male bodies and for muscularity as a discriminable dimension, extending a literature that has so far used female bodies, the body-size dimension, and reproduction methods (Alexi et al., 2018; Alexi et al., 2019; Turnbull et al., 2022). The small marginal attraction we observe is consistent with the attractive biases those studies reported. What the present method adds is a view of the opposing processes the marginal pattern conceals.

### Future directions

Several questions remain open for future work. The most immediate concerns the bodies and observers studied. The body serial-dependence literature, and much of the broader psychophysical and clinical work on body perception, has concentrated on female bodies, whereas our study used male bodies and male observers. The obvious next step is to measure serial dependence for female bodies with the same decomposition, connecting our trial-level results to the existing female-body literature. More informative still would be to cross the sex of the body with the sex of the observer, since the perceiver’s own body and lifetime visual experience may shape which dimensions are channel-coded. Brooks, Keen, et al. (2020) reported that body-composition aftereffects depend on the gender of both adapting and test bodies, which predicts that the fat-versus-muscularity dissociation we observe should itself vary with body sex and observer sex.

The framework is not specific to body composition. The same trial-level partialling, applied to continuously sampled stimuli, could ask which other socially meaningful dimensions are channel-coded and which are not. Candidates include other aspects of body perception and the wider domain of social vision: face identity, emotional expression, gender, ethnicity, and sex discrimination, several of which have already been shown to exhibit serial dependence in their marginal form (Liberman et al., 2014; Liberman et al., 2018; Taubert et al., 2016). None have yet been examined with the perceptual residual isolated from decisional carryover and its tuning profile estimated.

Perhaps the most important open question concerns the relationship between the trial-by-trial repulsion measured here and the prolonged-exposure aftereffects characterised by Brooks and colleagues. The two share a repulsive sign and, for body fat, a channel signature, which is suggestive of a common mechanism operating across timescales, but our data cannot establish this. Two approaches would help. (i) Within-subject correlations: if a single adaptation mechanism underlies both, the magnitude of an observer’s trial-by-trial repulsion should predict the magnitude of their prolonged-exposure aftereffect, whereas a dissociation would indicate at least partly separate processes. (ii) Systematic variation of stimulus duration: investigating how the perceptual and decisional components change as exposure increases from a single brief trial toward the minutes range of classical adaptation would show whether the trial-by-trial effect is the short-duration limit of the same continuous process or a qualitatively distinct one. Our finding that adaptation operates from a single preceding trial already implies that body perception recalibrates far more rapidly than the prolonged-adaptation paradigm suggests. Characterising the full duration function would establish whether one mechanism spans the whole range, and would speak directly to the neural question of whether the same body-selective populations support both.

### Conclusion

Serial dependence in male body discrimination, barely visible in the raw responses, resolves under trial-level partialling into a strong perceptual repulsion and a strong decisional attraction that ordinarily cancel. We show that both components, and the tuning profile of the perceptual one, can be obtained from a rapid two-alternative forced-choice task, and that the familiar narrow serial-dependence curve is the emergent sum of two broad opposing processes rather than a direct measure of channel width. The perceptual repulsion is more sharply tuned for body fat than for muscularity in our data, a difference that may reflect how readily channel tuning is resolved rather than its presence, and that links the moment-to-moment dynamics of body perception to the channel architecture previously inferred from prolonged adaptation.

## Declaration of generative AI and AI-assisted technologies in the manuscript preparation process

During the preparation of this work the authors used Claude (Anthropic) to improve the language and readability of the manuscript. After using this tool, the authors reviewed and edited the content as needed and take full responsibility for the content of the published article.

